# Multidimensional responses of ecological stability to eutrophication in grasslands

**DOI:** 10.1101/2023.04.16.537045

**Authors:** Qingqing Chen, Shaopeng Wang, Elizabeth T. Borer, Jonathan D. Bakker, Eric W. Seabloom, W. Stanley Harpole, Nico Eisenhauer, Ylva Lekberg, Yvonne M. Buckley, Jane A. Catford, Christiane Roscher, Ian Donohue, Sally A. Power, Pedro Daleo, Anne Ebeling, Johannes M. H. Knops, Jason P. Martina, Anu Eskelinen, John W. Morgan, Anita C. Risch, Maria C. Caldeira, Miguel N Bugalho, Risto Virtanen, Isabel C Barrio, Yujie Niu, Anke Jentsch, Carly J. Stevens, Juan Alberti, Yann Hautier

## Abstract

Eutrophication usually impacts biodiversity, species composition, and functioning of grassland communities. Whether such effects propagate to influence the stability of these community aspects is unknown. Using standardized experiments across 55 global grasslands, we quantified the effects of nutrient addition on five stability facets (i.e., temporal invariability and resistance during and recovery after dry and wet growing seasons) for three community aspects (i.e., aboveground biomass, community composition, and species richness). Nutrient addition reduced the temporal invariability and resistance of species richness and community composition, but not biomass, during dry and wet growing seasons. Temporal invariability and resistance during, but not recovery after, dry and wet growing seasons were strongly positively correlated in both ambient and eutrophic conditions. This indicates that maintaining and restoring the stability of plant communities requires increasing resistance rather than recovery. Harnessing the complexity of ecological stability provides new insights for grassland ecosystem sustainability in a changing world.

## Introduction

In 2020, the Convention on Biological Diversity reported that only 8% of the world’s nations met the target of limiting excess nutrients to a level that is not detrimental to ecosystem functions. This failure means that eutrophication, which disrupts diversity, functionality, and nature’s contributions to people^1^, could persist and pose a threat to our long-term survival and prosperity. Whether, and how, these changes propagate to affect ecosystem stability in the context of increasing climatic variability such as dry and wet climate extremes remain elusive.

In ecological studies, stability is a multifaceted concept that characterizes the ability of an ecosystem to minimize fluctuations in its properties against perturbations and variations in environmental conditions. Traditionally, stability has been assessed through temporal invariability (or temporal stability^2^), which involves calculating the mean of an ecosystem property divided by its deviation. Recent studies have emphasized the importance of measuring stability using resistance during and recovery from perturbations, such as droughts and floods^3,4^ (Fig. 1). However, most studies have focused on individual stability facets, such as temporal invariability or resistance to drought^5–7^. The few studies exploring the interrelationships among stability facets have yielded mixed results regarding the direction and strength of correlations^6,8–13^. For example, although increased resistance or recovery should lead to lower deviation from the long-term averages and higher temporal invariability, a meta-analysis suggests that higher resistance, not recovery, is the primary driver of temporal invariability^3^. These findings raise important questions about the correlations between stability facets and which facets are most critical for ecosystem stability.

**Fig. 1.**
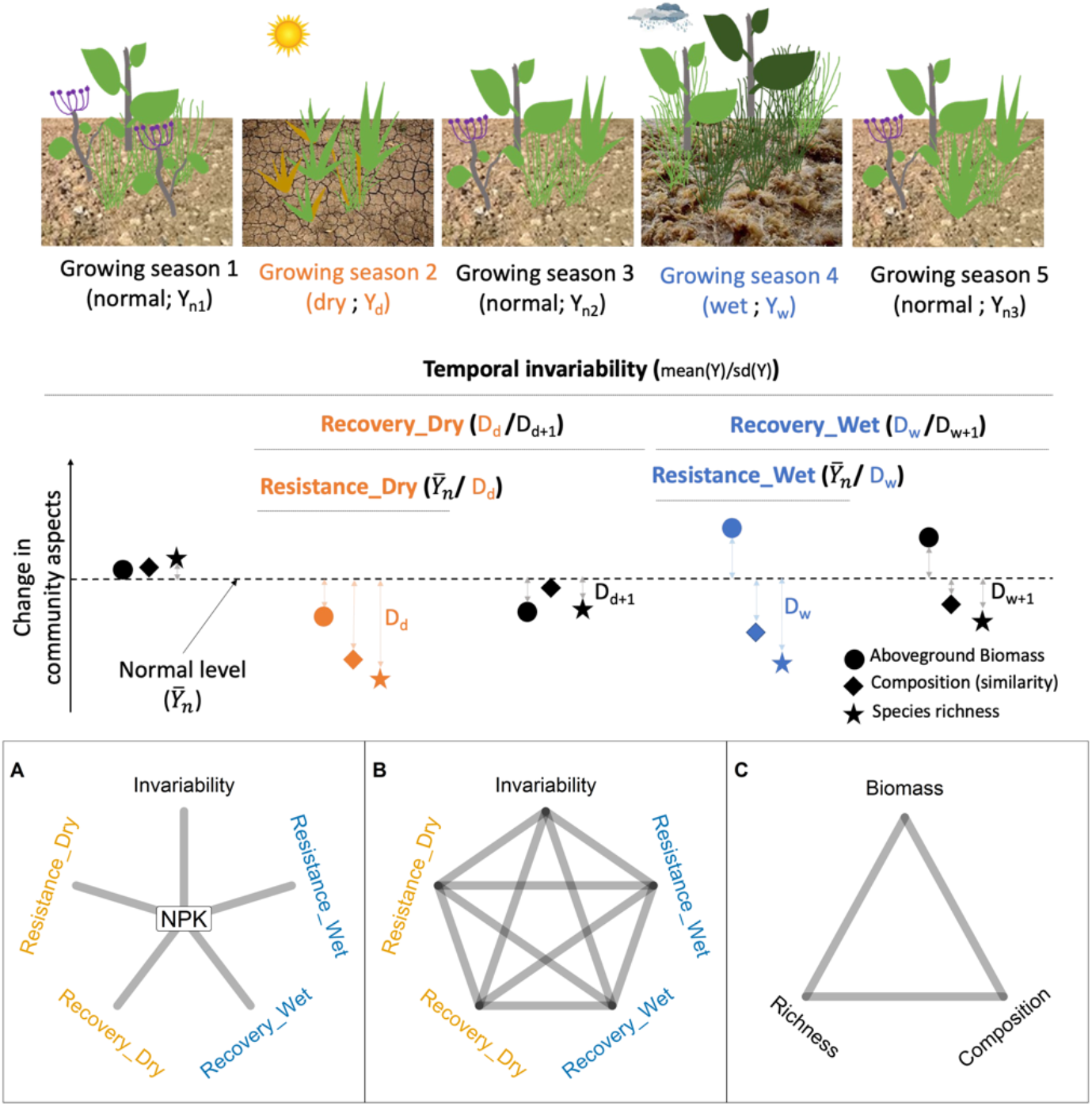
Graphical illustration of five stability facets in three community aspects investigated in this study. Methods used for quantifying stability facet are shown. We investigate the effects of nutrient addition on each stability facet (A), correlations between different stability facets within each community aspect (B), and correlations of stability between different community aspects for a given stability facet (C).

When stability facets are strongly correlated, they essentially represent a single dimension, which means that understanding the mechanisms behind one stability facet can provide crucial information for predicting and managing other facets. For example, if two stability facets are positively (resp. negatively) correlated, increasing ecological factors that boost one stability facet should lead to an increase (resp. decrease) in the other. However, if stability facets are negatively correlated, optimizing both facets becomes challenging because improving one will likely come at the expense of the other. On the other hand, when stability facets are not correlated, they effectively represent a high dimensionality, and the mechanisms that govern each stability facet are likely to be different.

Similarly, stability can be measured for multiple community aspects such as aboveground biomass, community composition, and species richness (Fig. 1) and the stability of different community aspects can correlate with one another^10,14,15^. While most studies on stability have focused on the stability of aboveground biomass, the stability of diversity aspects such as species richness and composition are essential for regulating ecosystem functions and maintaining ecosystem stability^16^. Understanding the various facets of stability in multiple community aspects simultaneously is crucial for predicting and managing ecosystem sustainability in the face of global environmental change^6^.

Previous studies in grassland ecosystems have shown that eutrophication usually reduces the temporal invariability of aboveground biomass^14^ and its resistance during dry climate extremes^17–19^. However, the impact of eutrophication on resistance during and recovery after wet climate extremes is relatively underexplored. Plant communities may exhibit asymmetrical responses during droughts and floods under eutrophic conditions, as both nutrients and water are crucial for plant survival and growth. Furthermore, eutrophication is known to decrease species richness and cause shifts in community composition which can in turn affect the temporal invariability of aboveground biomass^15^. However, whether eutrophication affects the stability of diversity aspects remains to be elucidated.

Additionally, eutrophication may alter the effective dimensionality of stability by changing the correlations among stability facets. For example, a greenhouse experiment showed that adding nutrients increases the correlations among different facets of stability, possibly due to enhanced interspecific competition and deterministic community assembly processes in eutrophic conditions^11,12^. However, eutrophication can also increase soil fertility and aboveground biomass, which may promote stochastic community assembly and weaken the association between different stability facets^14,20,21^. Thus, the net impacts of eutrophication on different stability facets and their correlations may reflect the combined effects of stochastic and deterministic processes in community dynamics. While some studies have shown that the positive correlation between temporal invariability in biomass and community composition remains strong under eutrophic conditions^11,14^, a systematic investigation of the correlations between different stability facets in various community aspects and their responses to eutrophication is still lacking. Understanding how environmental changes may impact the correlations between stability facets presents a new challenge to resolve the complexity of ecosystem sustainability and an opportunity to develop effective strategies for ecosystem management that maintain long-term sustainability.

Using 55 grassland sites spanning 5 continents with at least 4 years of standardized nutrient addition experiments, we tested whether nutrient addition alters five facets of stability measured for three community aspects (Fig. 1). The five stability facets include temporal invariability, resistance during and recovery after dry and wet growing seasons. The three community aspects include aboveground biomass, plant species richness, and plant community composition. In total, we investigated 15 stability facets. Moreover, we quantified whether and how nutrient addition alters relationships between different stability facets in different community aspects. We categorized dry, normal, and wet growing seasons for each site based on its historical standardized precipitation–evapotranspiration index (SPEI; dry: ≤ 25th percentile; wet: ≥75th percentile; normal: 25 - 75th percentile of standardized SPEI; Fig. S1-S5; Table S1-S2; see methods)^3^. In total, 150 dry, 247 normal, and 131 wet growing seasons were recorded across all sites during the study period.

## Results and discussion

First, we determined the impact of nutrient addition on five stability facets in three community aspects. Nutrient addition decreased temporal invariability and resistance in community composition and species richness during dry and wet growing seasons, but it did not affect stability in biomass (Fig. 2; Fig. S6; Fig. S7; Fig. S8; Table S3). Nutrient addition reduced resistance during dry and wet growing seasons in community composition, and resistance during dry growing seasons in species richness through the combined effects of reducing the normal levels and increasing deviation from the normal levels (Supplementary text; Fig. S7; Fig. S8). While nutrient addition reduced resistance during wet growing seasons in species richness primarily by reducing the normal levels of species richness (Supplementary text; Fig. S7). To determine whether nutrient addition had a stronger effect on rare species, we also calculated species diversity weighted by species cover. We found that nutrient addition decreased temporal invariability and resistance in cover-weighted species diversity in a similar way (Fig. S9). This suggests that sensitivity of plant diversity during dry and wet growing seasons under nutrient addition did not depend on rarity. While aboveground biomass was less sensitive than species richness and community composition, it could be attributed to species turnover compensating for biomass loss due to species loss^20^ or functional redundancy within communities sustaining a similar level of biomass even under species loss^22^. These results advance our understanding of ecosystem sustainability in changing environments by showing that nutrient addition increases the sensitivity of species richness and community composition during dry and wet growing seasons and decreased their temporal invariability.

**Fig. 2.**
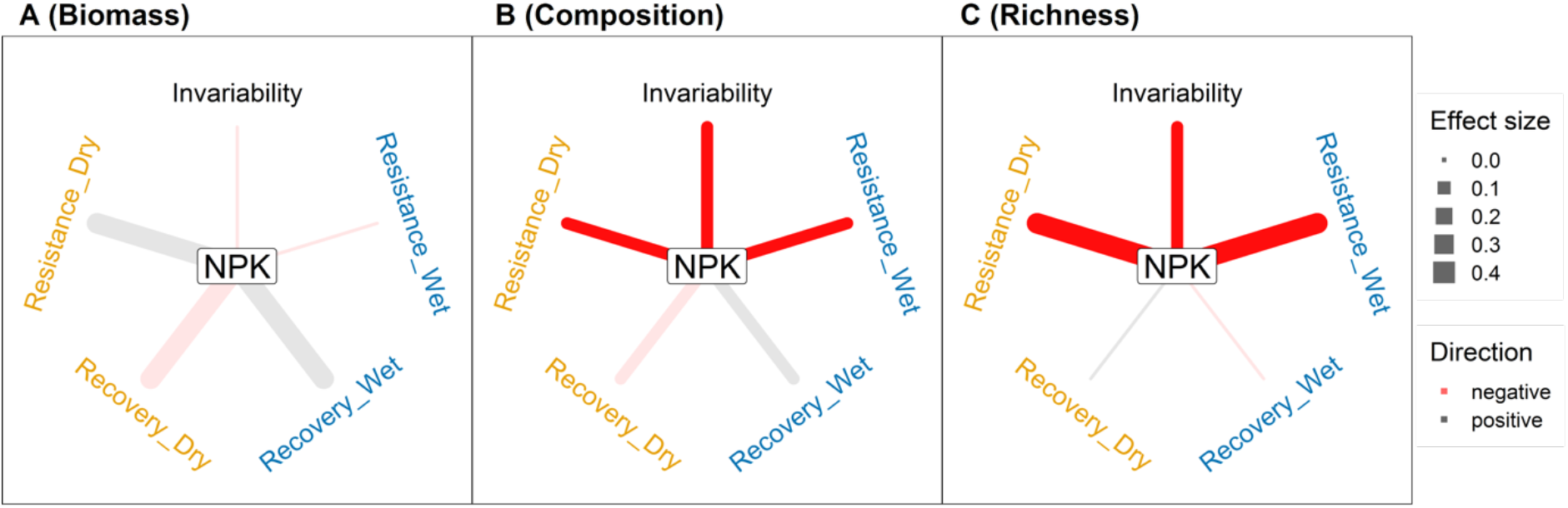
Effects of nutrient addition on each of the five stability facets in each of the three community aspects. Line width indicates effect size of nutrient addition on the respective stability facet. Saturated line colors represent significant effects at *p* ≤ 0.05, faded line colors represent non-significant effects. See Table S3 for test statistics and standard errors of the effect size.

Next, we analyzed the correlation among stability facets within each community aspect and their responses to nutrient addition. We calculated pairwise Pearson correlation coefficients among stability facets (resulting in 10 pairs from 5 facets) for each community aspect in either treatment at each site. Under ambient conditions, 4, 4, and 2 out of 10 pairs of stability facets were significantly correlated in biomass, community composition, and species richness, respectively. This suggests relatively high dimensionality of stability, in line with previous studies showing that one or two facets cannot adequately capture overall stability^6,10,11^. These relationships were generally similar under nutrient addition, although certain pairs of stability facets were weakened or enhanced (Fig. 3; Table S4). Specifically, nutrient addition weakened the negative correlation between resistance and recovery of biomass during dry growing seasons. Additionally, nutrient addition resulted in a negative correlation between temporal invariability and recovery after wet growing seasons and a positive correlation between resistance during dry and wet growing seasons in community composition, which were not correlated under ambient conditions. In both aboveground biomass and community composition, resistance during and recovery after dry or wet growing seasons were generally negatively correlated under both ambient and nutrient addition conditions. This indicates that plant communities that were more impacted during dry and wet growing seasons also recovered more, regardless of nutrient treatments. Such a trade-off between resistance and recovery is important for the maintenance of community composition and functions in the face of climate extremes. However, such a trade-off was not observed for species richness, implying that species richness may be more difficult to recover (from their normal levels) after perturbations. Notably, we found that temporal invariability was highly correlated with resistance, but not recovery, under both treatments in all three community aspects (Fig. 3). This is in line with findings from manipulated biodiversity experiments^3^ and demonstrates that long-term temporal invariability of community aspects relies on their short-term responses during climate extremes rather than their recovery afterwards. These correlations are robust to nutrient addition, indicating that strategies contributing to community resistance are also key for enhancing the temporal invariability of ecosystems in eutrophic conditions.

**Fig. 3.**
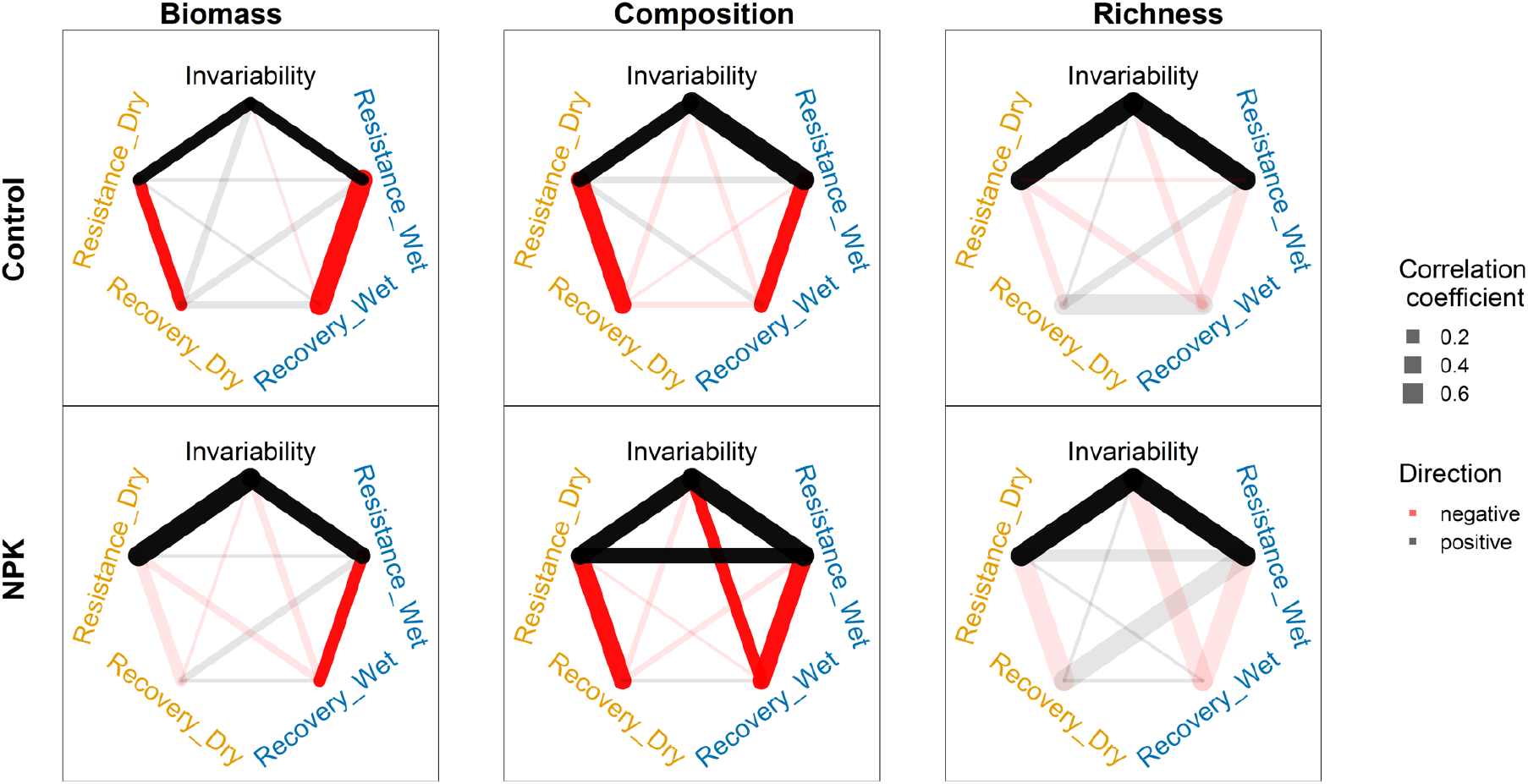
Pairwise correlations between five stability facets in three community aspects under ambient (control) and nutrient addition conditions. Pearson correlation coefficients were calculated between each pair of stability facets in each community aspect, averaged across sites under each treatment. Line width indicates the magnitude of correlation coefficients between two stability facets. Saturated line colors represent significant effects, corresponding to 95% confidence intervals that did not overlap with 0. See Table S4 for test statistics and 95% confidence intervals for each correlation coefficient.

Finally, we tested correlation among stability of community aspects for a given stability facet and their responses to nutrient addition. We found that the stability of the three community aspects were generally weakly correlated for each stability facet, under both ambient and nutrient addition conditions (Fig. 4; Table S5). This may be explained by the differential responses of biomass compared to community composition and species richness (Fig. 2). The correlations between stability of community composition and species richness were also weak possibly because community composition change was mainly driven by species replacement but not species loss^14^. Our findings differ from previous studies that focused on other organisms and single-site experiments, which showed that compositional stability and biomass stability are positively correlated^10,11^. This discrepancy may be due to previous studies mostly focusing on experimental pulse perturbations over short time frames and using methods that rely on stronger correlations among community aspects (e.g. using cover data for calculating functions). Overall, our results suggest that the stability of plant diversity and ecosystem functions represent separate dimensions and context-dependent strategies may be required for successful conservation efforts.

**Fig. 4.**
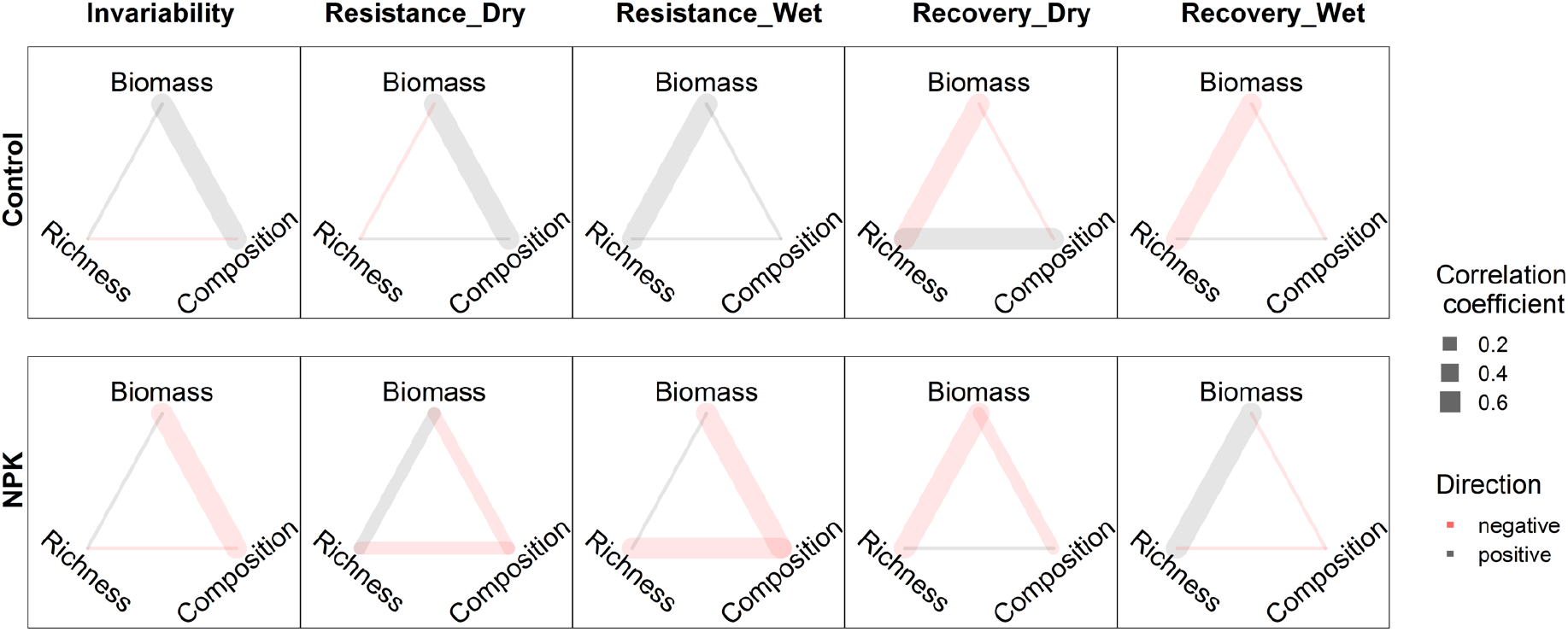
Pairwise correlations between stability of three community aspects for a given stability facet under ambient (control) and nutrient addition conditions. Pearson correlation coefficients were calculated between stability of each pair of community aspects, averaged across sites under each treatment. Line width indicates the magnitude of correlation coefficients between two stability facets. Saturated line colors represent significant effects, corresponding to 95% confidence intervals that did not overlap with 0. See Table S5 for test statistics and 95 % confidence intervals for each correlation coefficient.

Our study represents a globally coordinated efforts to investigate the multidimensional responses of ecological stability to eutrophication, by examining multiple facets of stability across multiple community aspects simultaneously. Our findings indicate that conserving plant diversity and community composition (e.g. pollinator plants or endangered species) may be more challenging than maintaining biomass production (e.g. agricultural grasslands) under climate extremes and eutrophication. Moreover, to maintain and restore temporal stability of plant communities, especially plant diversity, it is crucial to focus on increasing resistance rather than recovery. Our results reveal ecological stability is intrinsically multidimensional and that different facets of stability may be influenced by independent mechanisms and processes, underscoring the need to consider its multidimensional responses to environmental changes. By disentangle the concurrent impacts of global changes on ecosystems, our results provide new insights and opportunities to achieve a more holistic understanding of grassland ecosystem sustainability in a changing world.

## Methods

### Experimental Design

The study sites are part of the NutNet experiment^23,24^ (Fig. S1 and table S1). For the analyses here, we select plots assigned to one of two treatments: Control or Fertilized (NPK_+µ_). 5 m × 5 m plots were assigned to one of the two treatments in a randomized block design, typically with three blocks per site. NPK_+µ_ treatment plots were fertilized with nitrogen (N), phosphorus (P), and potassium with a combination of micronutrients and macronutrients (Fe, S, Mg, Mn, Cu, Zn, B, and Mo) as a one-time addition to the potassium treatment (K_+µ_). The micronutrient mix was applied once at the start of the experiment at a rate of 100 g m^-2^. N was supplied as time-release urea ((NH_2_)_2_CO), P was supplied as triple superphosphate (Ca(H_2_PO_4_)_2_), and K as potassium sulfate (K_2_SO_4_). N, P, and K were added annually at rates of 10 g m^-2^ y^-1^. Ammonium nitrate was used as the nitrogen source at some sites in 2007; however, urea was used in all subsequent years due to difficulties in procuring ammonium nitrate. An additional experiment at four NutNet sites shows that the effects of ammonium nitrate on plant richness and aboveground biomass were similar to that of urea^25^.

Data was retrieved in November 2022. The 55 sites included in this study met the following criteria: (1) plots were arranged in 3 blocks; (2) ≥ 4 years of post-treatment measurement; (3) during experimental years, at least one dry or wet growing season occurred (see “Defining climate extremes” for more details). These sites span five continents and include a wide range of grassland types. See Fig. S1, Fig. S2, and Table S1 for details of sites selected, experimental years, geolocation, and grassland types.

### Sampling protocol

All NutNet sites followed standard sampling protocols. A 1 ×1 m subplot within each plot was permanently marked for annual measurement of plant community composition. Species cover (%) was estimated visually for all species in the subplots; the total cover of living plants can exceed 100 % for multilayer canopies. Aboveground biomass was measured within the treatment plot, adjacent to the permanent subplot, by clipping all aboveground biomass within two 1 × 0.1 m strips (in total 0.2 m^2^), which were moved each year to avoid resampling the same location. For shrubs and subshrubs occurring in strips, we collected all leaves and current year’s woody growth. Biomass was dried at 60 °C (to constant mass) before weighing to the nearest 0.01 g, and expressed as g m^-2^. At most sites, cover was recorded once per year at peak biomass before fertilization. At some sites with strong seasonality, cover was recorded twice per year to include a complete list of species. For those sites, the maximum cover for each species and total biomass were used in the following analyses. The taxonomy was adjusted within sites to ensure consistent naming over time.

Specifically, when individuals could not be identified as species, they were aggregated at the genus level but referred to as “species” for simplicity.

### Defining climate extremes and stability facets for the three community aspects

Following ref ^3^, we used the standardized precipitation–evapotranspiration index (SPEI) to classify climate events for each site. SPEI was calculated as the standardized (z-score) water balance over the growing season each year (sum of precipitation – sum of evapotranspiration; mm) from 1901 to 2021. Precipitation and potential evapotranspiration used to calculate SPEI were downloaded from https://crudata.uea.ac.uk/cru/data/hrg/cru_ts_4.06^26^. At each site, we then classified the treatment years (growing seasons) into dry, normal, and wet using the cutoff of 0.67 or 1.28 sd (0.67 sd: occurring once every four years; 1.28 sd: occurring once per decade). In both cases, normal growing seasons were defined as -0.67 sd < SPEI < 0.67 sd. A cutoff of 0.67 sd (dry: ≤ 25th percentile; wet: ≥75th percentile) resulted in a much higher number of dry and wet growing seasons occurring at more sites (55 sites; Fig. S1; Table S1), which can increase the power of our statistical analyses. Results were similar when defining dry and wet growing seasons using SPEI 1.28 sd (dry: ≤ 10th percentile; wet: ≥90th percentile; 44 sites; resulting in 66 dry, 247 normal, and 58 wet growing seasons). Therefore, we present the results based on SPEI 0.67 sd in the main text and the results based on SPEI 1.28 sd in the supplementary file (Fig. S10-S12). To reduce confounding effects of dry and wet growing seasons occurring consecutively to resistance and recovery, we selected experimental years using the following three criteria. First, for resistance and recovery, if the previous non-normal growing season was not the same non-normal growing season, this growing season was eliminated. Second, for recovery, if the next growing season was a different non-normal growing season, this growing season was eliminated, the pre-treatment year and years with missing biomass were included for these two selection procedures. Third, when the two (or more) same non-normal growing seasons happen consecutively, recovery only calculated for the last growing season (must be followed by either a less but same non-normal growing season or a normal growing season). See table S2 for all combinations of three consecutive growing seasons and selection of the growing seasons for calculating resistance and recovery. To make clearer the selection of the years and calculation of resistance and recovery, we present changes in biomass in both control and nutrient addition treatments over years at all sites and at three blocks at site Look.us (Fig. S4 and Fig. S5).

Also, we summarized average changes in aboveground biomass, species richness, and community composition (from their normal levels) during and one year after dry and wet growing seasons in both treatments (Fig. S6-S8).

Methods used for quantifying stability facets in aboveground biomass and species richness are illustrated in Fig. 1. To remove variation due to directional change over time, we calculated temporal invariability in biomass and species richness after detrending. That is, we first used linear regression (function “lm”) to fit species richness or biomass against experimental years for each subplot, we then used the residuals from this model to calculate detrended standard deviation. Stability facets in biomass and species richness were log-transformed to improve homogeneity of variance. The composition-related facets of stability were calculated using Bray–Curtis dissimilarity metric based on cover data^27,28^. Temporal invariability was calculated as the overall community similarity over all experimental years (i.e., 1 - dissimilarity). The average cover for all species during normal growing seasons under each treatment within a block was constructed as a reference community for calculating resistance and recovery. Resistance was calculated as the similarity of the plant community under a non-normal growing season compared with the reference. Recovery was calculated as the ratio of similarity of the community one year after a non-normal growing season to that under a non-normal growing season. We used the function “beta.multi.abund” and “beta.pair.abund” from the R package betapart^27^ with Bray-Curtis dissimilarities to calculate temporal invariability, resistance and recovery, respectively. The data for temporal invariability and resistance, but not recovery, in community composition was bounded between -1 and 0. Resistance and recovery for all three community aspects were averaged over years to match the data structure of temporal invariability. For all stability facets in the three community aspects, higher values represent higher stability.

### Statistical analysis

All analyses were performed in R v.4.1.2^29^. We used linear mixed-effects models (function “lme”) from the R package “nlme” for the following analyses^30^. First, we tested whether nutrient addition impacted each stability facet for each community aspect. In the models, treatment (control vs. nutrient addition) was the fixed effect, while site and block nested within site were random effects. To look at whether rare species were more sensitive than common and dominant species during dry and wet growing seasons under nutrient addition, we looked at hill numbers with Q ranging from 0 to 2^31^. As the value of Q increases the abundant species weighs more in hill number. Second, we looked at the effects of nutrient addition on relationships among the five stability facets for each community aspect. We calculated Pearson correlation coefficients between every pair of stability facets for each treatment at each site, and then used the function “lme” to test the effect of treatment (fixed effect) on these correlation coefficients, with site as the random effect. Third, we looked at the effects of nutrient addition on relationships in stability among the three community aspects (for each stability facet). Similarly, we calculated Pearson correlation coefficients for a stability facet among all pairs of community aspects for each treatment at each site, and then used the function “lme” to test the effect of treatment (as fixed effects) on these correlation coefficients, with site as the random effect.

## Supporting information

supplementary file for Multidimensional responses of ecological stability to eutrophication in grasslands

## Author contributions (see Table S6 for more details)

Conceptualization: QC, SW, YH

Methodology and data analyses: QC, SW, YH, JB

R code checking: JB, YN

Investigation (data collection): all authors except QC and SW

Visualization: QC

Writing—original draft: QC, YH, SW

Writing—review & editing: all authors

## Acknowledgments

We thank researchers from the NutNet who contributed data to our analysis but are not listed as authors, supplementary Table S7 lists these researchers. We thank the Minnesota Supercomputing Institute for hosting project data and the Institute on the Environment for hosting Network meetings. Nitrogen fertilizer was donated to NutNet by Crop Production Services, Loveland, CO. This experiment is funded by individual researchers at the site scale.

## Data and materials availability

Data associated are available in Figshare (https://figshare.com/s/cc91715c26dfb827e0f4).

