## supplementary file for Multidimensional responses of ecological stability to eutrophication in grasslands for "Multidimensional responses of ecological stability to eutrophication in grasslands"

**Supplementary files**

To enable comparison among sites with varying growing conditions, we quantified resistance as the inverse of the proportional deviation of a community aspect from its normal levels during a dry or wet growing season. Thus, nutrient addition may decrease (or increase) resistance in a community aspect when it increases the magnitude of deviation more (or less) than it increases mean during normal growing seasons compared with the control.

Compared with the control, on average, nutrient addition led to a 48% increase in aboveground biomass during normal growing seasons (i.e. the normal levels). Nutrient addition decreased aboveground biomass during dry growing seasons while increasing it during wet growing seasons (i.e. large changes in biomass). This resulted in increased biomass deviation from the normal levels (note, change can be positive or negative values; while deviation is always positive values; Fig. S6). In contrast, nutrient addition decreased species richness by 19% during normal growing seasons. Nutrient addition had no effect on change in species richness during dry and wet seasons from the normal levels (Fig. S7). However, nutrient addition increased deviation in species richness during dry growing seasons, suggesting some sites increased while other sites decreased in species richness during dry growing seasons (relative to their normal levels). Nutrient addition did not impact deviation in species richness during wet growing seasons, suggesting most sites did not strongly change species richness during wet growing seasons (Fig. S7). Nutrient addition decreased community similarity during normal growing seasons, it also decreased community similarity during both dry and wet growing seasons relative to that of normal levels (Fig. S8).

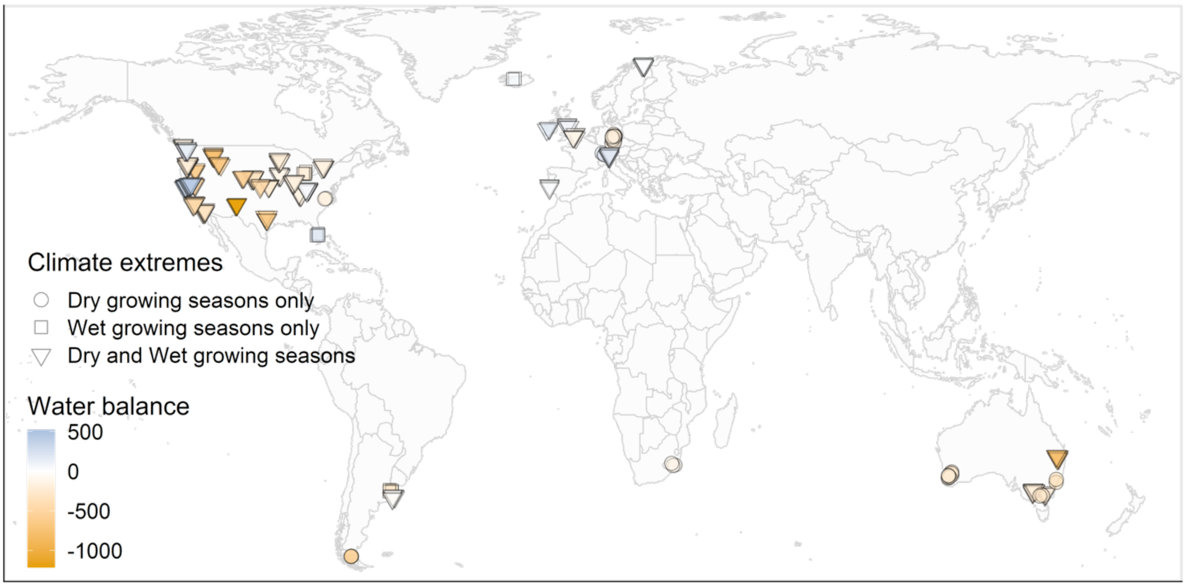


**Fig. S1. Sites used in this study, their geolocation, water balance, and climate extremes.** Water balance refers to the average values of precipitation minus evapotranspiration during growing seasons from 2007 to 2021. Dry/wet growing seasons only refer to sites where only dry or wet growing seasons were recorded during experimental years, dry and wet growing seasons refer to sites where both dry and wet growing seasons were recorded during experimental years. Sites are jittered to reduce overlap, see table S1 for more details.

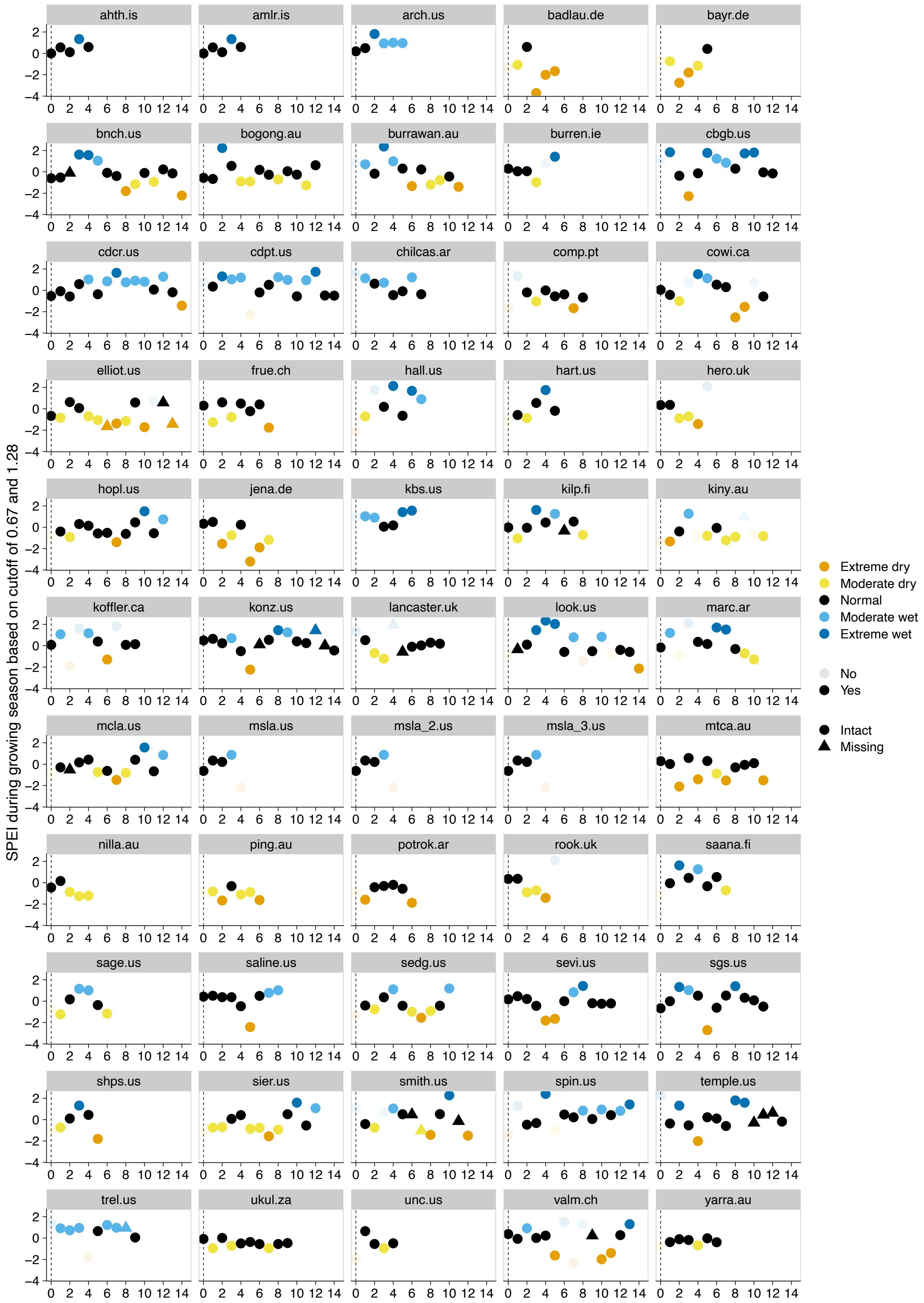


**Fig. S2. Extreme dry, moderate dry, normal, moderate wet, and extreme wet growing seasons at each site during the experimental years.** Positive and negative values indicate wetter and drier than normal growing seasons, respectively. SPEI accounts for water balance during the growing season based on data from January 1901 to 2021 at each site. To illustrate that dry and wet growing seasons depend on cutoffs used, at each site, we classified the treatment years (growing seasons) into extreme dry, moderate dry, normal, moderate wet, and extreme wet using the cutoff of 1.28 and 0.67 sd (1.28: occurring once per decade; 0.67: once every four years) following Isbell et al., (2015). That is, extreme dry: SPEI $\leq$ -1.28 sd; moderate dry: -1.28 sd < SPEI $\leq$ -0.67 sd; normal growing season: -0.67 sd < SPEI < 0.67 sd; moderate wet: 0.67 sd $\leq$ SPEI < 1.28 sd; and extreme wet: SPEI $\geq$ 1.28 sd. In analyses, we calculated resistance and recovery for dry and wet growing seasons based on either of the cutoffs. See table S1 for sites, the growing seasons, and experimental years used for the analyses.

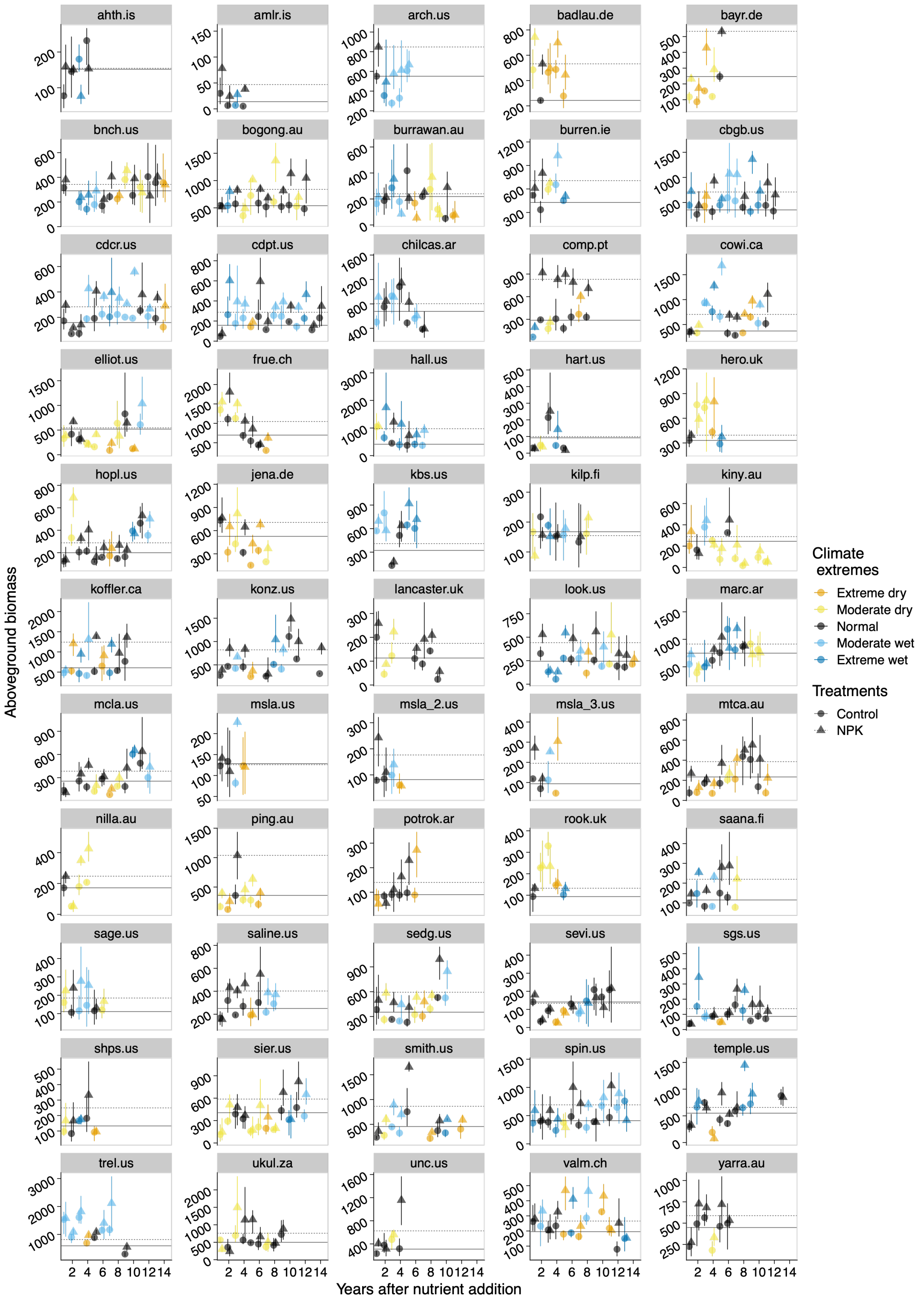


**Fig. S3. Aboveground biomass (g m$^{-2}$) over experimental years under ambient and nutrient addition conditions at each site.** Extreme dry, moderate dry, normal, moderate wet, and extreme wet using the cutoff of 1.28 and 0.67 sd (1.28: occurring once per decade; 0.67: once every four years) correspond to Fig. S2. Dots indicate raw live biomass data averaged over three blocks in each year at each site. Error bars are 95% bootstrapped confidence intervals. Solid and dashed lines represent average aboveground biomass in the control and nutrient addition conditions, respectively.

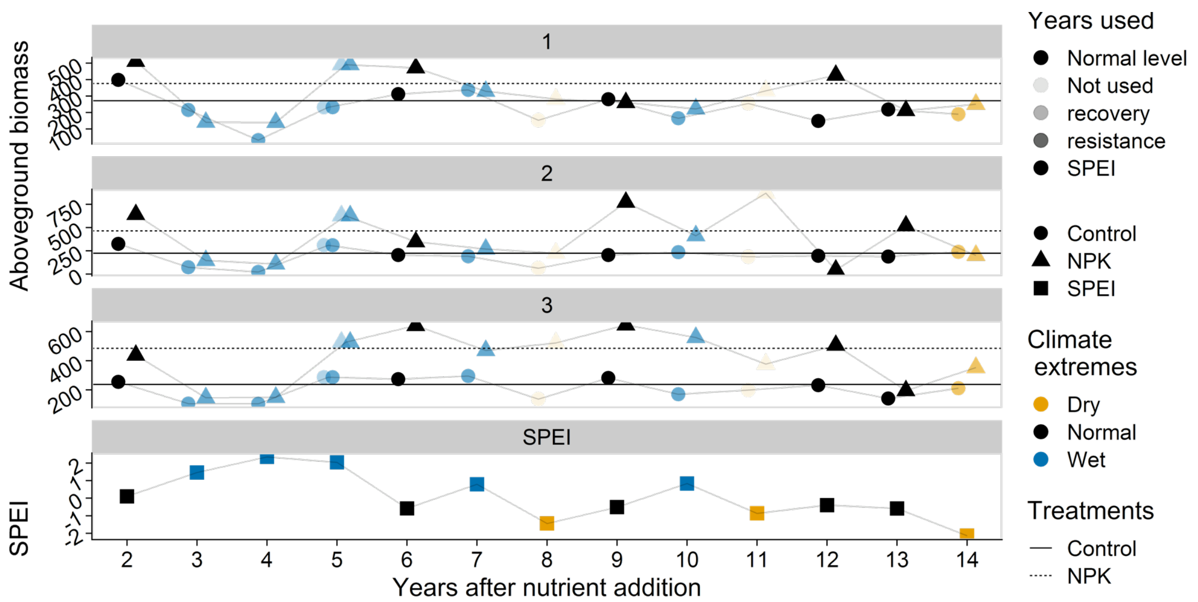


**Fig. S4. Aboveground biomass (g m$^{-2}$) over experimental years under ambient and nutrient addition conditions in three blocks (shown in three panels) at site Look.us.** Dry, normal, and wet growing seasons were categorized using the cutoff of 0.67 sd (i.e. a non-normal growing season occurred once every four years). Therefore, we considered both moderate and extreme dry growing seasons (from Fig. S2) as dry growing seasons, and moderate and extreme wet growing seasons (from Fig. S2) as wet growing seasons. This figure illustrates that the normal levels were calculated for each treatment in each block at each site. This figure also shows the years used for the calculating resistance and recovery.

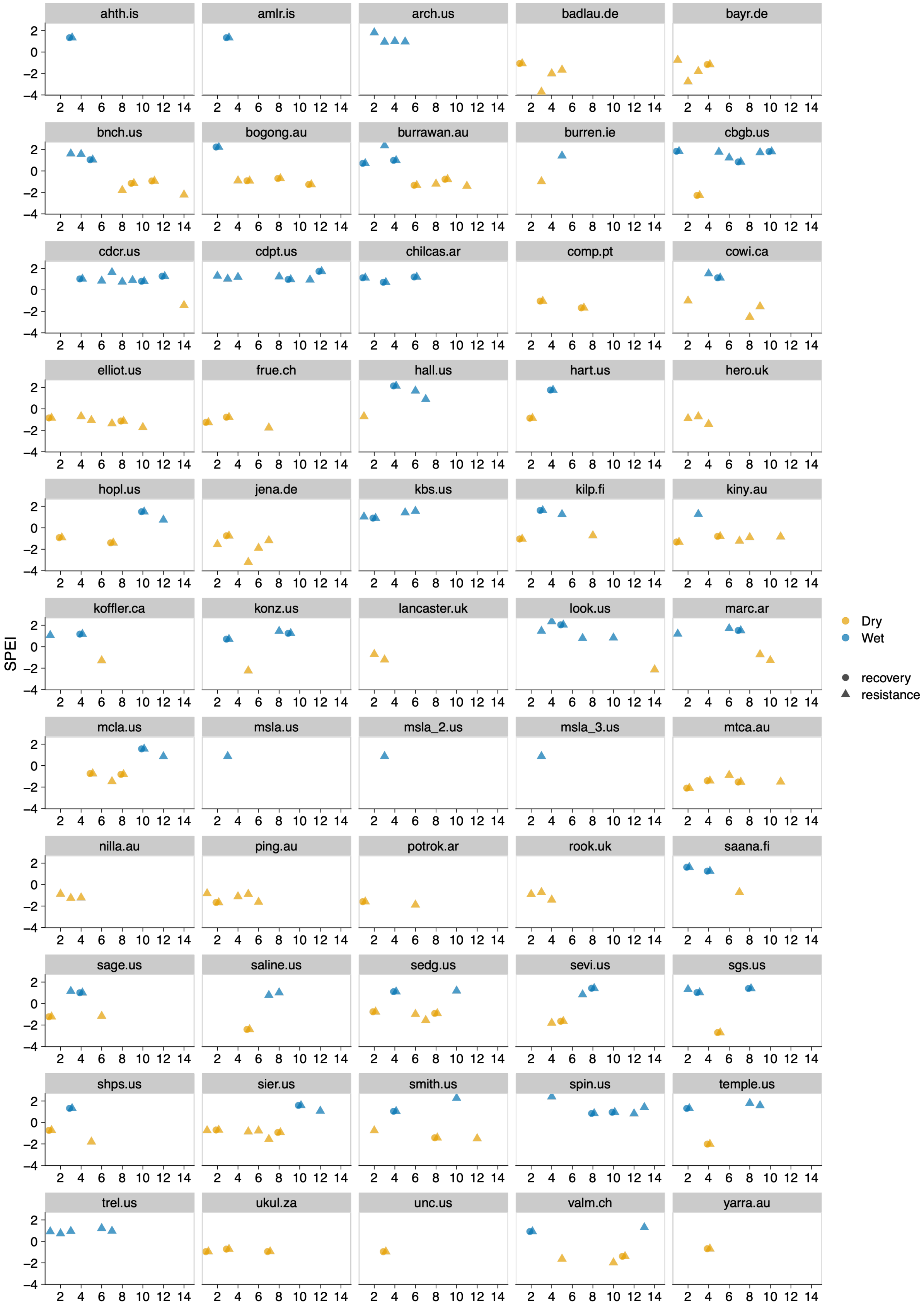


**Fig. S5. Dry and wet growing seasons at each site during the experimental years used for resistance and recovery.** Positive and negative values indicate wetter and drier than normal growing seasons, respectively. SPEI accounts for water balance during the growing season based on data from January 1901 to 2021 at each site. Dry and wet growing seasons are based on 25th and 75th percentiles of SPEI (cutoff of 0.67 sd). That is, we considered both moderate and extreme dry growing seasons from Fig. S2 as dry growing seasons, and moderate and extreme wet growing seasons as wet growing seasons. We also considered dry and wet growing seasons based on 10th and 90th percentiles of SPEI, where only extreme dry and wet growing seasons from Fig. S2 were used for calculating resistance and recovery, moderate dry and wet growing seasons were ignored. See Fig. S10 - Fig. S12 for results based on 10th and 90th percentiles of SPEI. See Table S2 for combinations of three consecutive growing seasons and selection of the growing seasons for calculating resistance and recovery.

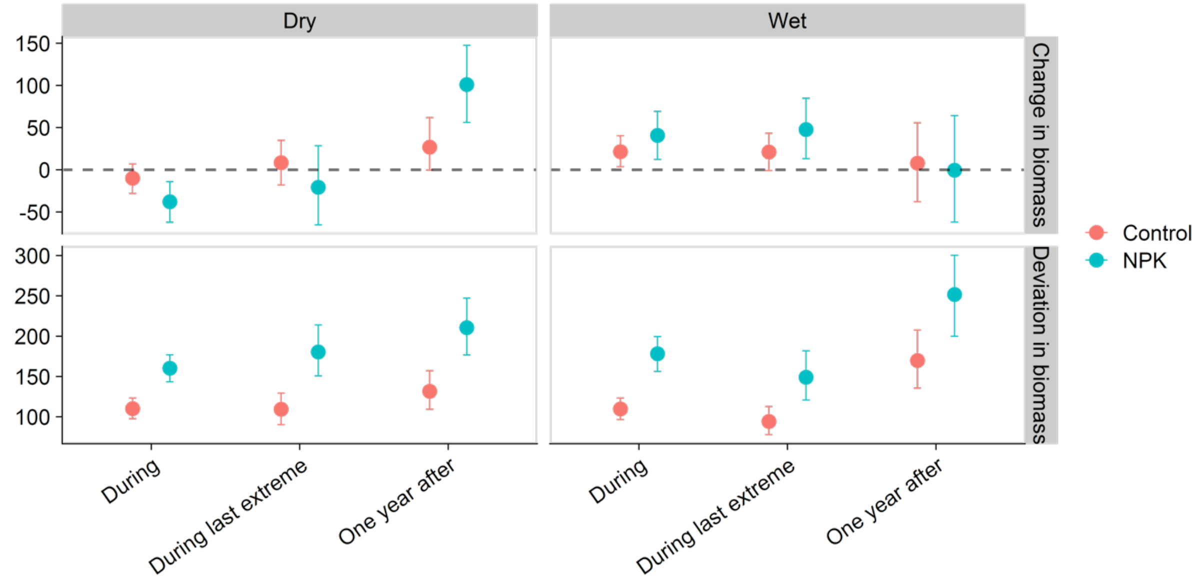


**Fig. S6. Change in aboveground biomass (g m$^{-2}$; upper panel) and the magnitude of biomass deviation from normal levels (lower panel) in control and nutrient addition treatments.** The normal levels are the means of aboveground biomass over normal growing seasons within treatments in each block at each site. During normal growing seasons, on average, nutrient addition increased aboveground biomass by 48% (control: 318.54; nutrient addition: 471.44; g.m$^{-2}$). Change in aboveground biomass refers to biomass differences from normal levels, thus values can be positive or negative. Deviation in aboveground biomass refers to absolute change in biomass from normal levels, thus values are positive only. “During the last extreme” refers to the last dry or wet growing season when more than one dry or wet growing seasons occur consecutively. “One year after” refers to a normal growing season after a dry or wet growing season. “During the last” and “one year after” were used to calculate recovery. Dots indicate average values over dry or wet growing seasons across 55 sites. Error bars are 95% bootstrapped confidence intervals.

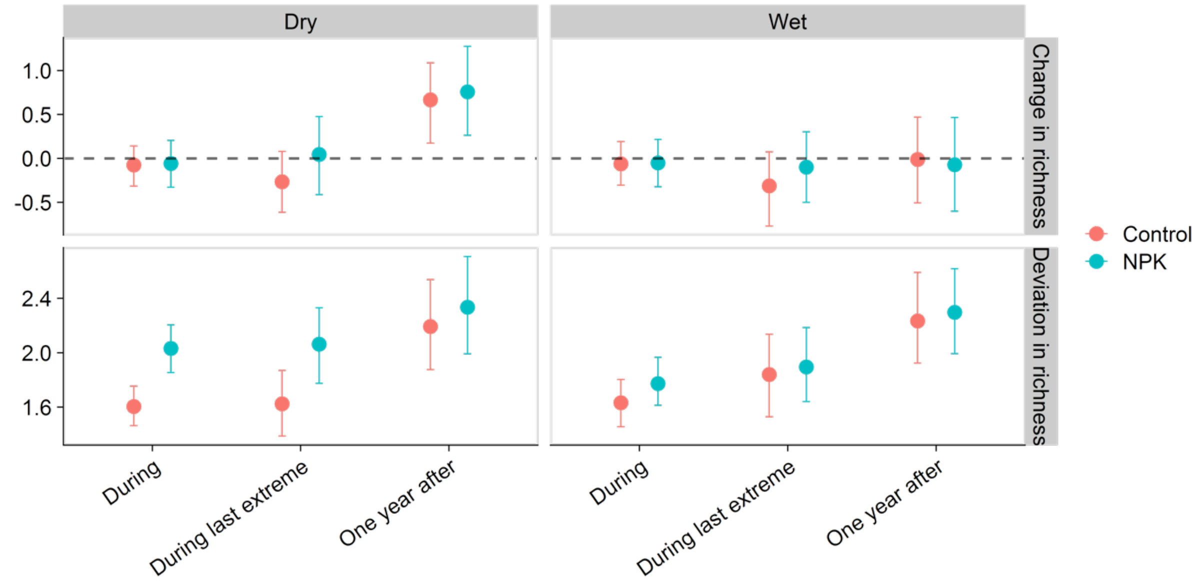


**Fig. S7. Change in species richness (spp m$^{-2}$; upper panel) and the magnitude of richness deviation from normal levels (lower panel) in control and nutrient addition plots during and one year after dry or wet growing seasons.** The normal levels are the means of species richness over normal growing seasons within treatments in each block at each site. During normal growing seasons, on average, nutrient addition decreased species richness by 19% (control: 12.18; nutrient addition: 9.82; spp.m$^{-2}$). Change in species richness refers to richness difference from normal levels, thus values can be positive or negative. Deviation in species richness refers to absolute change in richness from normal levels, thus values are positive only. “During the last extreme” refers to the last dry or wet growing season when more than one dry or wet growing seasons occur consecutively. “One year after” refers to a normal growing season after a dry or wet growing season. “During the last” and “one year after” were used to calculate recovery. Dots indicate average values during dry or wet growing seasons and across 55 sites. Error bars and thin lines are 95% bootstrapped confidence intervals.

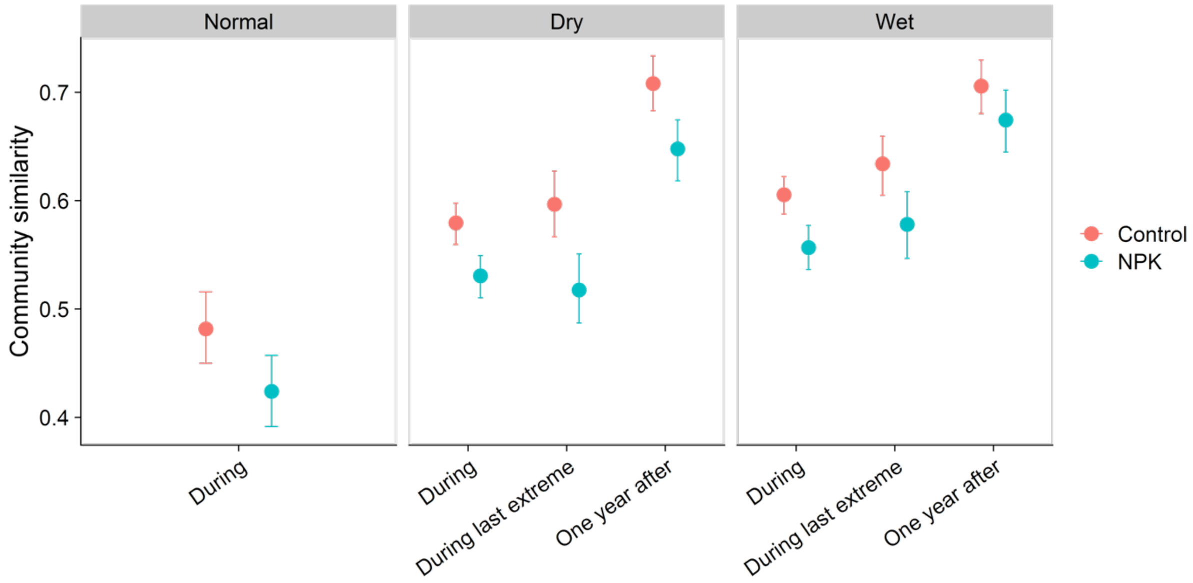


**Fig. S8. Average community similarity during normal growing seasons, during and one year after dry and wet growing seasons.** During the last refers to the last dry or wet growing season when more than one dry or wet growing seasons occur consecutively. “During the last extreme” refers to the last dry or wet growing season when more than one dry or wet growing seasons occur consecutively. “One year after” refers to a normal growing season after a dry or wet growing season. “During the last” and “one year after” were used to calculate recovery. Dots are means, error bars are 95% bootstrapped confidence intervals.

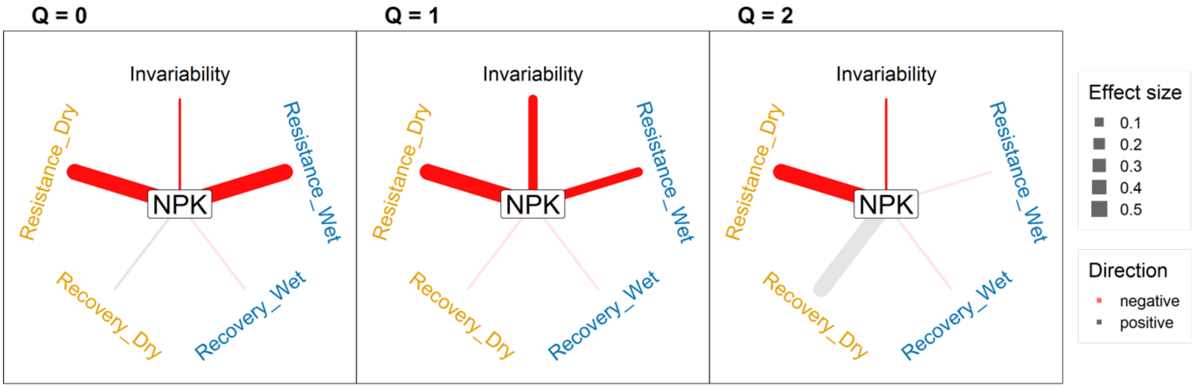


**Fig. S9. Effects of nutrient addition on five facets stability of species diversity quantified with hill numbers (with varying Q).** As the value of Q increased, the abundant species weighed more in this diversity index. Saturated line colors represent significant effects at p ≤ 0.05, faded line colors represent non-significant effects.

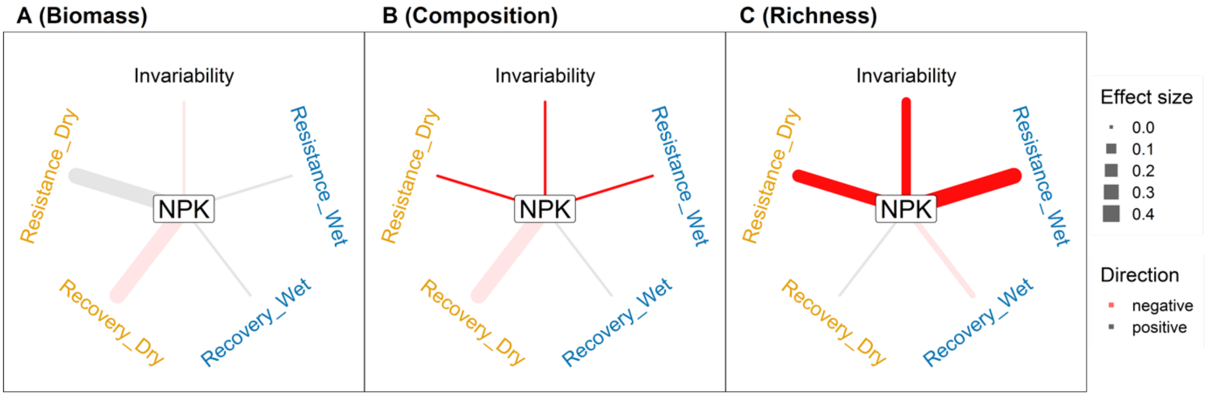


**Fig. S10. Effects of nutrient addition on each of the five stability facets in each of the three community three aspects.** Compare with Figure 2 which used less strict definitions (25th and 75th percentiles of SPEI) of dry and wet growing seasons but more sites, here dry and wet growing seasons based on 10th and 90th percentiles of SPEI. Saturated line colors represent significant effects at $p \leq 0.05$, faded line colors represent non-significant effects.

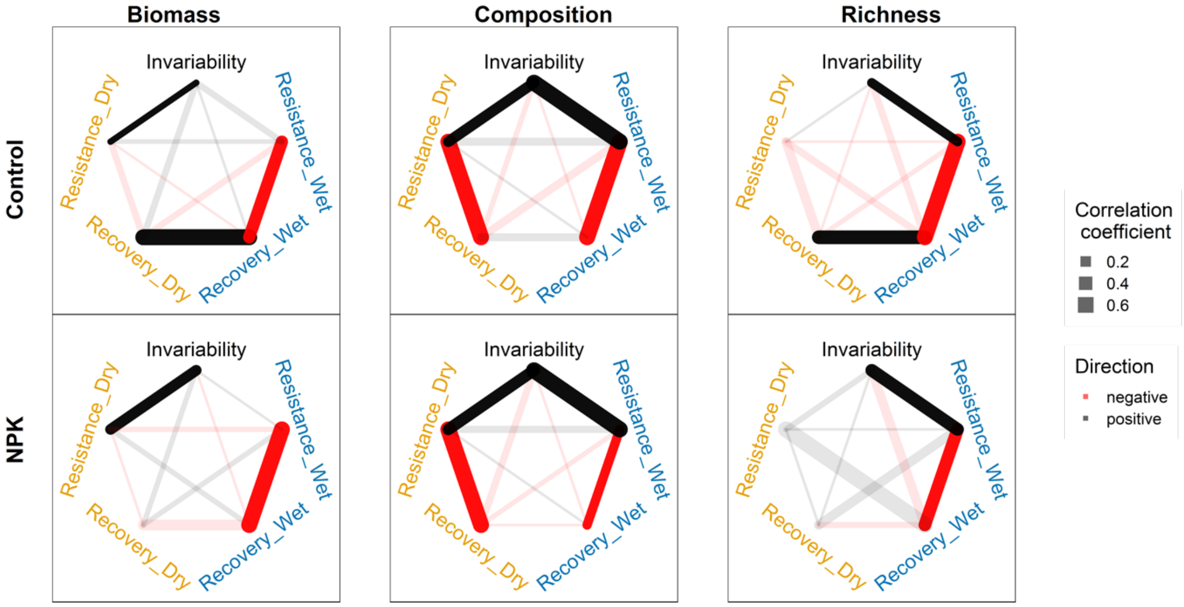


**Fig. S11. Pairwise correlations among five stability facets in three community aspects under ambient and nutrient addition conditions.** Compare with Figure 3 which used less strict definitions (25th and 75th percentiles of SPEI) of dry and wet growing seasons but more sites, here dry and wet growing seasons based on 10th and 90th percentiles of SPEI. The significant effects (saturated colors) correspond to 95 % confidence intervals of a correlation coefficient does not overlap with 0.

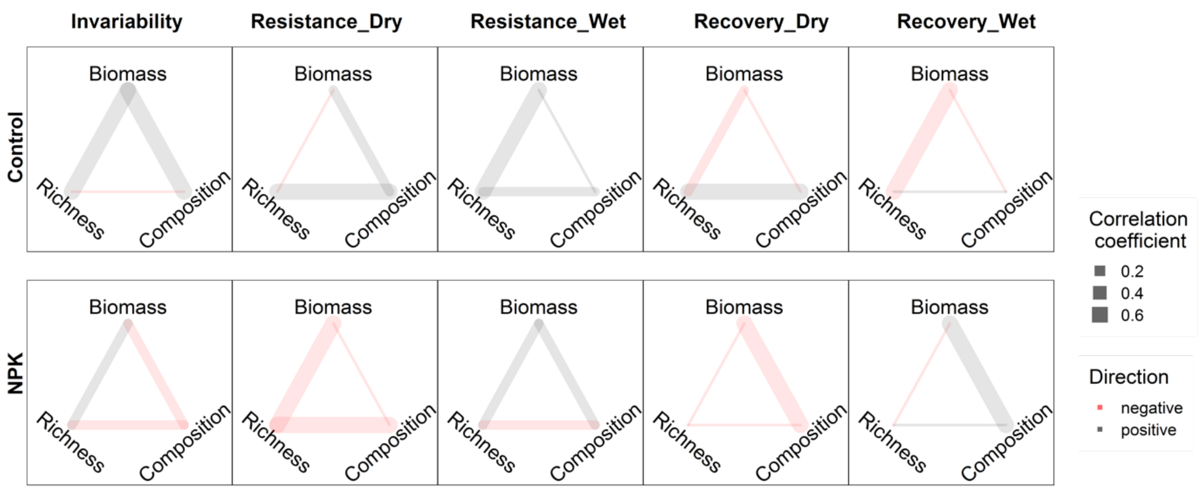


**Fig. S12. Pairwise correlations of stability among three community aspects under ambient and nutrient addition conditions.** Compare with Figure 4 which used less strict definitions (25th and 75th percentiles of SPEI) of dry and wet growing seasons but more sites, here dry and wet growing seasons based on 10th and 90th percentiles of SPEI. The significant effects (saturated colors) correspond to 95 % confidence intervals of a correlation coefficient does not overlap with 0.

**Table S1. Information for 55 sites included in the analyses.** Water balance refers to the average values of precipitation minus evapotranspiration during the growing seasons from 2007 to 2021. Growing seasons were estimated by site PIs, start and end growing season is the month when vegetation turns green and vegetation turns brown.

| site_code | Habitat | Continent | Latitude | Longitude | Growing season (start to end) | Water balance (2007-2021) |
|---|---|---|---|---|---|---|
| ahth.is | heathland | Europe | 65.13 | -19.67 | 6-9 | 57.72 |
| amlr.is | desert grassland | Europe | 65.13 | -19.67 | 6-9 | 57.72 |
| arch.us | mixedgrass prairie | North America | 27.17 | -81.22 | 5-10 | 177.28 |
| badlau.de | old field | Europe | 51.39 | 11.88 | 4-10 | -263.08 |
| bayr.de | mesic grassland | Europe | 49.92 | 11.58 | 3-9 | -202.68 |
| bnch.us | montane grassland | North America | 44.28 | -121.97 | 4-8 | -410.42 |
| bogong.au | alpine grassland | Australia | -36.87 | 147.25 | 10-1 | -161.27 |
| burrawan.au | semiarid grassland | Australia | -27.74 | 151.14 | 10-5 | -768.43 |
| burren.ie | calcareous grassland | Europe | 53.07 | -8.99 | 2-8 | 169.66 |
| cbgb.us | tallgrass prairie | North America | 41.79 | -93.39 | 5-10 | -121.70 |
| cdcr.us | tallgrass prairie | North America | 45.42 | -93.21 | 4-8 | -198.09 |
| cdpt.us | shortgrass prairie | North America | 41.21 | -101.64 | 4-7 | -320.66 |
| chilcas.ar | mesic grassland | South America | -36.28 | -58.27 | 8-3 | -283.02 |
| comp.pt | annual grassland | Europe | 38.83 | -8.79 | 10-5 | 26.15 |
| cowi.ca | old field | North America | 48.81 | -123.63 | 4-7 | -126.39 |
| elliot.us | annual grassland | North America | 32.88 | -117.05 | 11-4 | -320.77 |
| frue.ch | pasture | Europe | 47.11 | 8.54 | 4-9 | 200.73 |
| hall.us | tallgrass prairie | North America | 36.87 | -86.70 | 4-9 | -146.23 |
| hart.us | shrub steppe | North America | 42.72 | -119.50 | 10-7 | -663.66 |
| hero.uk | mesic grassland | Europe | 51.41 | -0.64 | 4-10 | -114.17 |
| hopl.us | annual grassland | North America | 39.01 | -123.06 | 11-4 | 519.56 |
| jena.de | grassland | Europe | 50.94 | 11.53 | 3-10 | -179.13 |
| kbs.us | old field | North America | 42.41 | -85.39 | 4-9 | -198.20 |
| kilp.fi | tundra grassland | Europe | 69.06 | 20.87 | 6-9 | 24.72 |
| kiny.au | semiarid grassland | Australia | -36.20 | 143.75 | 5-10 | -201.76 |
| koffler.ca | pasture | North America | 44.02 | -79.54 | 4-8 | -160.54 |

| site_code | Habitat | Continent | Latitude | Longitude | Growing season (start to end) | Water balance (2007-2021) |
|---|---|---|---|---|---|---|
| konz.us | tallgrass prairie | North America | 39.07 | -96.58 | 5-9 | -233.79 |
| lancaster.uk | mesic grassland | Europe | 53.99 | -2.63 | 3-8 | 76.77 |
| look.us | montane grassland | North America | 44.21 | -122.13 | 3-8 | -205.99 |
| marc.ar | grassland | South America | -37.72 | -57.42 | 4-12 | -39.35 |
| mcla.us | annual grassland | North America | 38.86 | -122.41 | 11-4 | 271.61 |
| msla.us | grassland | North America | 46.66 | -114.00 | 4-7 | -389.77 |
| msla_2.us | grassland | North America | 46.66 | -114.00 | 4-7 | -779.53 |
| msla_3.us | grassland | North America | 46.66 | -114.00 | 4-7 | -779.53 |
| mtca.au | savanna | Australia | -31.78 | 117.61 | 8-10 | -261.25 |
| nilla.au | old field | Australia | -36.90 | 146.01 | 2-1 | -288.90 |
| ping.au | old field | Australia | -32.50 | 116.97 | 4-10 | -230.83 |
| potrok.ar | semiarid grassland | South America | -51.92 | -70.41 | 10-4 | -581.11 |
| rook.uk | mesic grassland | Europe | 51.41 | -0.64 | 4-10 | -114.17 |
| saana.fi | montane grassland | Europe | 69.04 | 20.84 | 6-9 | 24.72 |
| sage.us | montane grassland | North America | 39.43 | -120.24 | 4-7 | -466.61 |
| saline.us | mixedgrass prairie | North America | 39.05 | -99.10 | 5-9 | -482.76 |
| sedg.us | annual grassland | North America | 34.70 | -120.02 | 11-7 | -459.86 |
| sevi.us | desert grassland | North America | 34.36 | -106.69 | 4-11 | -1214.75 |
| sgs.us | shortgrass prairie | North America | 40.82 | -104.77 | 4-8 | -565.40 |
| shps.us | shrub steppe | North America | 44.26 | -112.21 | 4-9 | -698.95 |
| sier.us | annual grassland | North America | 39.24 | -121.28 | 11-4 | 415.05 |
| smith.us | mesic grassland | North America | 48.21 | -122.62 | 10-6 | 130.83 |
| spin.us | pasture | North America | 38.13 | -84.50 | 3-5 | 47.75 |
| temple.us | tallgrass prairie | North America | 31.04 | -97.35 | 3-10 | -656.61 |
| trel.us | tallgrass prairie | North America | 40.08 | -88.83 | 4-9 | -166.75 |
| ukul.za | mesic grassland | Africa | -29.67 | 30.40 | 9-4 | -118.91 |
| unc.us | old field | North America | 36.01 | -79.02 | 4-9 | -195.07 |
| valm.ch | alpine grassland | Europe | 46.63 | 10.37 | 6-8 | 176.95 |

| site_code | Habitat | Continent | Latitude | Longitude | Growing season (start to end) | Water balance (2007-2021) |
|---|---|---|---|---|---|---|
| yarra.au | mesic grassland | Australia | -33.61 | 150.74 | 9-3 | -407.76 |

**Table S2. Combinations of three consecutive growing seasons and selection of the growing seasons for calculating resistance and recovery.**

| One year before | A given dry or wet growing season | One year after | Year for resistance | Year for recovery |
|---|---|---|---|---|
| Dry | Dry | Dry | Y | N |
| Dry | Dry | Normal | Y | Y |
| Dry | Dry | Wet | Y | N |
| Normal | Dry | Dry | Y | N |
| Normal | Dry | Normal | Y | Y |
| Normal | Dry | Wet | Y | N |
| Wet | Dry | Dry | N | N |
| Wet | Dry | Normal | N | N |
| Wet | Dry | Wet | N | N |
| Dry | Wet | Dry | N | N |
| Dry | Wet | Normal | N | N |
| Dry | Wet | Wet | N | N |
| Normal | Wet | Dry | Y | N |
| Normal | Wet | Normal | Y | Y |
| Normal | Wet | Wet | Y | N |
| Wet | Wet | Dry | Y | N |
| Wet | Wet | Normal | Y | Y |
| Wet | Wet | Wet | Y | N |

**Table S3. Model summaries for the fixed effects of nutrient addition on the five stability facets in three community aspects.** Models were specified as lme (stability facet ~ Treatment, random=~1|site/block). Stability facets based on aboveground biomass, species richness, but not community composition, are on the log scale. r2m (marginal R2): proportion of variance explained by the fixed effects in the model; r2c (conditional R2): proportion of variance explained by the fixed and random effects. SD (block); standard deviation for the random effect of blocks (nested within sites); SD (site): standard deviation for the random effect of sites.

| Community aspect | Sability facet | Terms | Value | Std Error | DF | t-value | p | r2m | r2c | SD (block) | SD (site) |
|---|---|---|---|---|---|---|---|---|---|---|---|
| biomass | invariability | (Intercept) | 0.94 | 0.05 | 164 | 18.97 | 0.00 | 0.00 | 0.47 | 0.10 | 0.30 |
| biomass | invariability | trtNPK | -0.02 | 0.04 | 164 | -0.54 | 0.59 | 0.00 | 0.47 | 0.10 | 0.30 |
| biomass | recovery_Dry | (Intercept) | 0.67 | 0.18 | 83 | 3.70 | 0.00 | 0.00 | 0.21 | 0.00 | 0.64 |
| biomass | recovery_Dry | trtNPK | -0.13 | 0.19 | 83 | -0.69 | 0.49 | 0.00 | 0.21 | 0.00 | 0.64 |
| biomass | recovery_Wet | (Intercept) | 0.32 | 0.15 | 92 | 2.11 | 0.04 | 0.00 | 0.21 | 0.00 | 0.56 |
| biomass | recovery_Wet | trtNPK | 0.14 | 0.16 | 92 | 0.85 | 0.39 | 0.00 | 0.21 | 0.00 | 0.56 |
| biomass | resistance_Dry | (Intercept) | 1.16 | 0.09 | 131 | 12.37 | 0.00 | 0.00 | 0.11 | 0.00 | 0.32 |
| biomass | resistance_Dry | trtNPK | 0.10 | 0.11 | 131 | 0.87 | 0.39 | 0.00 | 0.11 | 0.00 | 0.32 |
| biomass | resistance_Wet | (Intercept) | 1.32 | 0.12 | 116 | 10.69 | 0.00 | 0.00 | 0.32 | 0.00 | 0.59 |
| biomass | resistance_Wet | trtNPK | -0.02 | 0.11 | 116 | -0.19 | 0.85 | 0.00 | 0.32 | 0.00 | 0.59 |
| composition | recovery_Dry | (Intercept) | 1.43 | 0.13 | 92 | 10.92 | 0.00 | 0.00 | 0.21 | 0.12 | 0.48 |
| composition | recovery_Dry | trtNPK | -0.01 | 0.14 | 92 | -0.10 | 0.92 | 0.00 | 0.21 | 0.12 | 0.48 |

| Community aspect | Sability facet | Terms | Value | Std Error | DF | t-value | p | r2m | r2c | SD (block) | SD (site) |
|---|---|---|---|---|---|---|---|---|---|---|---|
| composition | recovery_Wet | (Intercept) | 1.23 | 0.09 | 92 | 13.93 | 0.00 | 0.00 | 0.19 | 0.18 | 0.28 |
| composition | recovery_Wet | trtNPK | 0.04 | 0.10 | 92 | 0.39 | 0.70 | 0.00 | 0.19 | 0.18 | 0.28 |
| composition | resistance_Dry | (Intercept) | 0.59 | 0.02 | 131 | 28.06 | 0.00 | 0.03 | 0.60 | 0.00 | 0.13 |
| composition | resistance_Dry | trtNPK | -0.06 | 0.01 | 131 | -4.30 | 0.00 | 0.03 | 0.60 | 0.00 | 0.13 |
| composition | resistance_Wet | (Intercept) | 0.61 | 0.02 | 116 | 25.83 | 0.00 | 0.02 | 0.63 | 0.00 | 0.13 |
| composition | resistance_Wet | trtNPK | -0.05 | 0.01 | 116 | -3.63 | 0.00 | 0.02 | 0.63 | 0.00 | 0.13 |
| richness | invariability | (Intercept) | 1.88 | 0.07 | 164 | 26.78 | 0.00 | 0.02 | 0.62 | 0.00 | 0.47 |
| richness | invariability | trtNPK | -0.19 | 0.04 | 164 | -4.47 | 0.00 | 0.02 | 0.62 | 0.00 | 0.47 |
| richness | recovery_Dry | (Intercept) | 0.32 | 0.14 | 74 | 2.24 | 0.03 | 0.00 | 0.21 | 0.00 | 0.49 |
| richness | recovery_Dry | trtNPK | 0.08 | 0.15 | 74 | 0.54 | 0.59 | 0.00 | 0.21 | 0.00 | 0.49 |
| richness | recovery_Wet | (Intercept) | 0.24 | 0.13 | 80 | 1.81 | 0.07 | 0.00 | 0.22 | 0.00 | 0.48 |
| richness | recovery_Wet | trtNPK | -0.07 | 0.14 | 80 | -0.50 | 0.62 | 0.00 | 0.22 | 0.00 | 0.48 |
| richness | resistance_Dry | (Intercept) | 2.04 | 0.11 | 127 | 18.39 | 0.00 | 0.02 | 0.50 | 0.07 | 0.63 |
| richness | resistance_Dry | trtNPK | -0.28 | 0.08 | 127 | -3.50 | 0.00 | 0.02 | 0.50 | 0.07 | 0.63 |
| richness | resistance_Wet | (Intercept) | 2.01 | 0.10 | 109 | 20.32 | 0.00 | 0.02 | 0.38 | 0.09 | 0.49 |
| richness | resistance_Wet | trtNPK | -0.22 | 0.09 | 109 | -2.53 | 0.01 | 0.02 | 0.38 | 0.09 | 0.49 |

**Table S4. Model summaries for the fixed effects of nutrient addition on pairwise correlations among the five stability facets in each community aspect.** Models were specified as lme (pairwise correlations ~ Treatment, random=~1|site). Stability facets based on aboveground biomass, species richness, but not community composition, are on the log scale. r2m (marginal R2): proportion of variance explained by the fixed effects in the model; r2c (conditional R2): proportion of variance explained by the fixed and random effects. SD (site): standard deviation for the random effect of sites.

| Community aspect | Correlation type | Treatment | Emmean | df | lower.CL | upper.CL | r2m | r2c | SD (site) |
|---|---|---|---|---|---|---|---|---|---|
| biomass | inv_recov.d | Control | 0.12 | 27 | -0.18 | 0.42 | 0.01 | 0.01 | 0.00 |
| biomass | inv_recov.d | NPK | -0.05 | 27 | -0.35 | 0.25 | 0.01 | 0.01 | 0.00 |
| biomass | inv_recov.w | Control | -0.04 | 30 | -0.31 | 0.23 | 0.00 | 0.09 | 0.22 |
| biomass | inv_recov.w | NPK | -0.14 | 30 | -0.41 | 0.13 | 0.00 | 0.09 | 0.22 |
| biomass | inv_resis.d | Control | 0.29 | 43 | 0.08 | 0.49 | 0.01 | 0.01 | 0.00 |
| biomass | inv_resis.d | NPK | 0.41 | 43 | 0.20 | 0.61 | 0.01 | 0.01 | 0.00 |
| biomass | inv_resis.w | Control | 0.26 | 38 | 0.03 | 0.48 | 0.00 | 0.00 | 0.00 |
| biomass | inv_resis.w | NPK | 0.35 | 38 | 0.12 | 0.57 | 0.00 | 0.00 | 0.00 |
| biomass | recov.d_recov.w | Control | 0.15 | 16 | -0.24 | 0.55 | 0.00 | 0.00 | 0.00 |
| biomass | recov.d_recov.w | NPK | 0.10 | 16 | -0.30 | 0.49 | 0.00 | 0.00 | 0.00 |
| biomass | resis.d_recov.d | Control | -0.29 | 27 | -0.58 | 0.00 | 0.00 | 0.16 | 0.29 |
| biomass | resis.d_recov.d | NPK | -0.25 | 27 | -0.54 | 0.04 | 0.00 | 0.16 | 0.29 |
| biomass | resis.d_recov.w | Control | 0.03 | 24 | -0.26 | 0.31 | 0.02 | 0.02 | 0.00 |
| biomass | resis.d_recov.w | NPK | -0.18 | 24 | -0.47 | 0.10 | 0.02 | 0.02 | 0.00 |
| biomass | resis.d_resis.w | Control | 0.06 | 27 | -0.23 | 0.35 | 0.00 | 0.18 | 0.32 |
| biomass | resis.d_resis.w | NPK | 0.05 | 27 | -0.24 | 0.34 | 0.00 | 0.18 | 0.32 |
| biomass | resis.w_recov.d | Control | 0.15 | 18 | -0.23 | 0.54 | 0.00 | 0.06 | 0.20 |
| biomass | resis.w_recov.d | NPK | 0.11 | 18 | -0.28 | 0.49 | 0.00 | 0.06 | 0.20 |
| biomass | resis.w_recov.w | Control | -0.48 | 30 | -0.72 | -0.24 | 0.02 | 0.13 | 0.22 |
| biomass | resis.w_recov.w | NPK | -0.28 | 30 | -0.52 | -0.04 | 0.02 | 0.13 | 0.22 |
| composition | inv_recov.d | Control | -0.17 | 30 | -0.43 | 0.09 | 0.00 | 0.00 | 0.00 |
| composition | inv_recov.d | NPK | -0.14 | 30 | -0.40 | 0.12 | 0.00 | 0.00 | 0.00 |
| composition | inv_recov.w | Control | -0.14 | 30 | -0.38 | 0.09 | 0.02 | 0.02 | 0.00 |
| composition | inv_recov.w | NPK | -0.32 | 30 | -0.56 | -0.09 | 0.02 | 0.02 | 0.00 |
| composition | inv_resis.d | Control | 0.47 | 43 | 0.27 | 0.66 | 0.00 | 0.10 | 0.20 |
| composition | inv_resis.d | NPK | 0.54 | 43 | 0.35 | 0.74 | 0.00 | 0.10 | 0.20 |
| composition | inv_resis.w | Control | 0.64 | 38 | 0.46 | 0.82 | 0.00 | 0.04 | 0.12 |
| composition | inv_resis.w | NPK | 0.63 | 38 | 0.44 | 0.81 | 0.00 | 0.04 | 0.12 |
| composition | recov.d_recov.w | Control | -0.17 | 16 | -0.56 | 0.22 | 0.03 | 0.03 | 0.00 |
| composition | recov.d_recov.w | NPK | 0.09 | 16 | -0.30 | 0.48 | 0.03 | 0.03 | 0.00 |
| composition | resis.d_recov.d | Control | -0.51 | 30 | -0.74 | -0.29 | 0.00 | 0.00 | 0.00 |

| Community aspect | Correlation type | Treatment | Emmean | df | lower.CL | upper.CL | r2m | r2c | SD (site) |
|---|---|---|---|---|---|---|---|---|---|
| composition | resis.d_recov.d | NPK | -0.51 | 30 | -0.74 | -0.29 | 0.00 | 0.00 | 0.00 |
| composition | resis.d_recov.w | Control | 0.11 | 24 | -0.19 | 0.42 | 0.01 | 0.20 | 0.32 |
| composition | resis.d_recov.w | NPK | -0.07 | 24 | -0.37 | 0.24 | 0.01 | 0.20 | 0.32 |
| composition | resis.d_resis.w | Control | 0.14 | 27 | -0.14 | 0.42 | 0.03 | 0.14 | 0.24 |
| composition | resis.d_resis.w | NPK | 0.41 | 27 | 0.13 | 0.68 | 0.03 | 0.14 | 0.24 |
| composition | resis.w_recov.d | Control | -0.04 | 18 | -0.39 | 0.32 | 0.00 | 0.00 | 0.00 |
| composition | resis.w_recov.d | NPK | -0.12 | 18 | -0.48 | 0.23 | 0.00 | 0.00 | 0.00 |
| composition | resis.w_recov.w | Control | -0.42 | 30 | -0.66 | -0.17 | 0.01 | 0.08 | 0.18 |
| composition | resis.w_recov.w | NPK | -0.52 | 30 | -0.76 | -0.28 | 0.01 | 0.08 | 0.18 |
| richness | inv_recov.d | Control | 0.07 | 21 | -0.25 | 0.39 | 0.00 | 0.13 | 0.26 |
| richness | inv_recov.d | NPK | 0.06 | 21 | -0.25 | 0.38 | 0.00 | 0.13 | 0.26 |
| richness | inv_recov.w | Control | -0.20 | 24 | -0.47 | 0.07 | 0.00 | 0.23 | 0.31 |
| richness | inv_recov.w | NPK | -0.20 | 24 | -0.47 | 0.07 | 0.00 | 0.23 | 0.31 |
| richness | inv_resis.d | Control | 0.31 | 39 | 0.08 | 0.53 | 0.00 | 0.00 | 0.00 |
| richness | inv_resis.d | NPK | 0.27 | 39 | 0.05 | 0.50 | 0.00 | 0.00 | 0.00 |
| richness | inv_resis.w | Control | 0.31 | 32 | 0.06 | 0.56 | 0.00 | 0.24 | 0.34 |
| richness | inv_resis.w | NPK | 0.28 | 32 | 0.03 | 0.53 | 0.00 | 0.24 | 0.34 |
| richness | recov.d_recov.w | Control | 0.32 | 14 | -0.06 | 0.70 | 0.05 | 0.05 | 0.00 |
| richness | recov.d_recov.w | NPK | 0.03 | 14 | -0.35 | 0.40 | 0.05 | 0.05 | 0.00 |
| richness | resis.d_recov.d | Control | -0.16 | 19 | -0.49 | 0.17 | 0.01 | 0.01 | 0.00 |
| richness | resis.d_recov.d | NPK | -0.28 | 19 | -0.61 | 0.05 | 0.01 | 0.01 | 0.00 |
| richness | resis.d_recov.w | Control | -0.20 | 18 | -0.51 | 0.12 | 0.04 | 0.06 | 0.10 |
| richness | resis.d_recov.w | NPK | 0.06 | 18 | -0.26 | 0.37 | 0.04 | 0.06 | 0.10 |
| richness | resis.d_resis.w | Control | -0.01 | 22 | -0.32 | 0.30 | 0.02 | 0.02 | 0.00 |
| richness | resis.d_resis.w | NPK | 0.17 | 22 | -0.14 | 0.48 | 0.02 | 0.02 | 0.00 |
| richness | resis.w_recov.d | Control | 0.12 | 15 | -0.29 | 0.53 | 0.00 | 0.10 | 0.24 |
| richness | resis.w_recov.d | NPK | 0.21 | 15 | -0.20 | 0.62 | 0.00 | 0.10 | 0.24 |
| richness | resis.w_recov.w | Control | -0.22 | 21 | -0.52 | 0.08 | 0.00 | 0.00 | 0.00 |
| richness | resis.w_recov.w | NPK | -0.23 | 21 | -0.53 | 0.07 | 0.00 | 0.00 | 0.00 |

**Table S5. Model summaries for the fixed effects of nutrient addition on pairwise correlations of stability among the three community aspects.** Models were specified as lme (pairwise correlations ~ Treatment, random=~1|site). Stability facets based on aboveground biomass, species richness, but not community composition, are on the log scale. r2m (marginal R2): proportion of variance explained by the fixed effects in the model; r2c (conditional R2): proportion of variance explained by the fixed and random effects. SD (site): standard deviation for the random effect of sites.

| Stability facet | Correlation type | Treatment | Emmean | df | lower.CL | upper.CL | r2m | r2c | SD (site) |
|---|---|---|---|---|---|---|---|---|---|
| invariability | bio_com | Control | 0.10 | 54 | -0.09 | 0.29 | 0.03 | 0.03 | 0.00 |
| invariability | bio_com | NPK | -0.12 | 54 | -0.31 | 0.06 | 0.03 | 0.03 | 0.00 |
| invariability | bio_div | Control | 0.07 | 54 | -0.12 | 0.26 | 0.00 | 0.00 | 0.00 |
| invariability | bio_div | NPK | 0.09 | 54 | -0.10 | 0.29 | 0.00 | 0.00 | 0.00 |
| invariability | com_div | Control | -0.03 | 54 | -0.23 | 0.16 | 0.00 | 0.11 | 0.24 |
| invariability | com_div | NPK | -0.08 | 54 | -0.28 | 0.11 | 0.00 | 0.11 | 0.24 |
| recovery_Dry | bio_com | Control | -0.02 | 27 | -0.29 | 0.26 | 0.01 | 0.01 | 0.00 |
| recovery_Dry | bio_com | NPK | -0.18 | 27 | -0.46 | 0.09 | 0.01 | 0.01 | 0.00 |
| recovery_Dry | bio_div | Control | -0.25 | 21 | -0.58 | 0.08 | 0.00 | 0.00 | 0.00 |
| recovery_Dry | bio_div | NPK | -0.26 | 21 | -0.59 | 0.07 | 0.00 | 0.00 | 0.00 |
| recovery_Dry | com_div | Control | 0.29 | 21 | -0.02 | 0.61 | 0.04 | 0.04 | 0.00 |
| recovery_Dry | com_div | NPK | 0.01 | 21 | -0.30 | 0.33 | 0.04 | 0.04 | 0.00 |
| recovery_Wet | bio_com | Control | -0.08 | 30 | -0.35 | 0.18 | 0.00 | 0.11 | 0.24 |
| recovery_Wet | bio_com | NPK | -0.01 | 30 | -0.28 | 0.25 | 0.00 | 0.11 | 0.24 |
| recovery_Wet | bio_div | Control | -0.13 | 24 | -0.40 | 0.15 | 0.03 | 0.36 | 0.39 |
| recovery_Wet | bio_div | NPK | 0.11 | 24 | -0.16 | 0.38 | 0.03 | 0.36 | 0.39 |
| recovery_Wet | com_div | Control | 0.04 | 24 | -0.27 | 0.34 | 0.00 | 0.00 | 0.00 |
| recovery_Wet | com_div | NPK | -0.05 | 24 | -0.36 | 0.25 | 0.00 | 0.00 | 0.00 |

| Stability facet | Correlation type | Treatment | Emmean | df | lower.CL | upper.CL | r2m | r2c | SD (site) |
|---|---|---|---|---|---|---|---|---|---|
| resistance_Dry | bio_com | Control | 0.15 | 43 | -0.07 | 0.37 | 0.02 | 0.09 | 0.19 |
| resistance_Dry | bio_com | NPK | -0.05 | 43 | -0.26 | 0.17 | 0.02 | 0.09 | 0.19 |
| resistance_Dry | bio_div | Control | -0.09 | 39 | -0.33 | 0.14 | 0.01 | 0.01 | 0.00 |
| resistance_Dry | bio_div | NPK | 0.03 | 39 | -0.20 | 0.27 | 0.01 | 0.01 | 0.00 |
| resistance_Dry | com_div | Control | 0.08 | 39 | -0.14 | 0.30 | 0.01 | 0.01 | 0.00 |
| resistance_Dry | com_div | NPK | -0.05 | 39 | -0.27 | 0.17 | 0.01 | 0.01 | 0.00 |
| resistance_Wet | bio_com | Control | 0.03 | 38 | -0.20 | 0.27 | 0.01 | 0.01 | 0.00 |
| resistance_Wet | bio_com | NPK | -0.13 | 38 | -0.36 | 0.10 | 0.01 | 0.01 | 0.00 |
| resistance_Wet | bio_div | Control | 0.14 | 32 | -0.11 | 0.40 | 0.01 | 0.01 | 0.00 |
| resistance_Wet | bio_div | NPK | 0.02 | 32 | -0.24 | 0.27 | 0.01 | 0.01 | 0.00 |
| resistance_Wet | com_div | Control | 0.00 | 32 | -0.26 | 0.25 | 0.00 | 0.00 | 0.00 |
| resistance_Wet | com_div | NPK | -0.10 | 32 | -0.35 | 0.15 | 0.00 | 0.00 | 0.00 |

**Table S6. Contributions of each author to the manuscript.**

| Full name | Site(s) used in analysis | Developed and framed research question(s) | Analyzed data | Contributed to data analyses | Wrote the paper | Contributed to paper writing | Site coordinator | Nutrient Network coordinator | Site-level acknowledgments (funding, access, etc) |
|---|---|---|---|---|---|---|---|---|---|
| Qingqing Chen | NA | x | x | NA | x | NA | NA | NA | NA |
| Yann Hautier | Frue.ch | x | x | NA | NA | x | x | NA | NA |
| Shaopeng Wang | NA | x | x | NA | NA | x | NA | NA | NA |
| Johannes M H Knops | cdpt | NA | NA | NA | NA | x | x | NA | NA |
| Anita C. Risch | valm.ch | NA | NA | NA | NA | x | x | NA | NA |
| Eric W. Seabloom | bnch.us, cdcr.us, hopl.us, look.us, mcla.us, sier.us | NA | NA | NA | NA | x | x | x | NA |
| Anne Ebeling | jena.de | NA | NA | NA | NA | x | x | NA | NA |
| John W. Morgan | bogong.au, kiny.au | NA | NA | NA | NA | x | x | NA | NA |
| Christiane Roscher | jena.de | NA | NA | NA | NA | x | x | NA | NA |
| Sally A. Power | NA | NA | NA | NA | NA | x | x | NA | NA |
| Jason P. Martina | temple.us | NA | NA | NA | NA | x | x | NA | NA |
| Jane A. Catford | nilla.au | NA | NA | NA | NA | X | X | NA | NA |
| Maria C. Caldeira | comp.pt | NA | NA | NA | NA | x | x | NA | Portuguese Science Foundation (FCT) for funding the research unit CEF |

| Full name | Site(s) used in analysis | Developed and framed research question(s) | Analyzed data | Contributed to data analyses | Wrote the paper | Contributed to paper writing | Site coordinator | Nutrient Network coordinator | Site-level acknowledgements (funding, access, etc) |
|---|---|---|---|---|---|---|---|---|---|
| | | | | | | | | | (UIDB/00239/2020) and to Companhia das Lezirias for field access |
| Miguel N Bugalho | comp.pt | NA | NA | NA | NA | x | x | NA | Portuguese Science Foundation (FCT) for funding the research unit CEABN-InBIO (UIDB/50027/2020) and to Rui Alves for logistic suport and granting access to study site |
| Elizabeth T Borer | bnch.us, cdcr.us, hopl.us, look.us, mcla.us, sier.us | NA | NA | NA | NA | x | x | x | NA |
| Risto Virtanen | kilp.fi, saana.fi | NA | NA | NA | NA | x | x | NA | NA |
| Anu Eskelinen | kilp.fi, saana.fi | NA | NA | NA | NA | x | x | NA | NA |
| Nico Eisenhauer | badlau.de | NA | NA | NA | NA | x | x | NA | Further support came from the German Centre for Integrative Biodiversity Research (iDiv) Halle-Jena-Leipzig, funded by the German Research Foundation (FZT 118, 202548816). |
| W Stanley Harpole | mcla.us, sier.us | NA | NA | NA | NA | x | x | NA | Further support came from the German Centre for Integrative Biodiversity Research (iDiv) Halle-Jena-Leipzig, funded by the German Research Foundation (FZT 118, 202548816). |

| Full name | Site(s) used in analysis | Developed and framed research question(s) | Analyzed data | Contributed to data analyses | Wrote the paper | Contributed to paper writing | Site coordinator | Nutrient Network coordinator | Site-level acknowledgments (funding, access, etc) |
|---|---|---|---|---|---|---|---|---|---|
| Ian Donohue | burren.ie | NA | NA | NA | NA | x | x | NA | National Parks and Wildlife Service, Ireland |
| Yvonne M. Buckley | burren.ie | NA | NA | NA | NA | x | x | NA | National Parks and Wildlife Service, Ireland |
| Isabel C Barrio | ahth.is, amlr.is | NA | NA | NA | NA | x | x | NA | University of Iceland Research Fund (2015), Soil Conservation Service of Iceland, Orkurannsóknasjóður Landsvirkjunnar (NÝR-09-2017, NÝR-14-2018, NÝR-12-2019) |
| Jonathan D. Bakker | smith.us | NA | NA | x | NA | x | x | NA | NA |
| Anke Jentsch | bayr.de | NA | NA | NA | NA | x | x | NA | Federal Ministry of Education and Research BMBF (FKZ 031B0516C, 031B1067C) |
| Yujie Niu | bayr.de | NA | NA | double checking the code | NA | x | NA | NA | Y. Niu is a Humboldt fellow funded by Alexander von Humboldt-Stiftung. |
| Juan Alberti | marc.ar | NA | NA | x | NA | x | NA | NA | NA |
| Pedro Daleo | marc.ar | NA | NA | x | NA | x | NA | NA | NA |
| Carly J. Stevens | lancaster.uk | NA | NA | NA | NA | x | x | NA | NA |
| Ylva Lekberg | msla_us; msla_2us; msla_3us | NA | NA | NA | NA | x | x | NA | MPG Ranch |

**Table S7. Principal investigators who contribute data are not authors; site names match those in Table S1.** Their effort in providing data is critical to this manuscript.

| site_code | PI.name | Institution |
|---|---|---|
| arch.us | Elizabeth Boughton | MacArthur Agro-ecology Research Center |
| badlau.de | Sylvia Haider | Martin-Luther-UniversitÃ¤t Halle-Wittenberg |
| badlau.de | Julia Siebert | German Centre for Integrative Biodiversity Research (iDiv) |
| bayr.de | Marie Spohn | NULL |

| site_code | PI.name | Institution |
|---|---|---|
| bogong.au | Joslin Moore | University of Melbourne |
| burrawan.au | Jennifer Firn | Queensland University of Technology |
| cbgb.us | Lori Biederman | Iowa State University |
| cbgb.us | Kirsten Hofmockel | Iowa State University |
| cbgb.us | Lauren Sullivan | Iowa State University |
| cdcr.us | Adam Kay | University of St. Thomas |
| cdpt.us | Johannes Knops | University of Nebraska, Lincoln |
| chilcas.ar | Enrique Chaneton | Universidad de Buenos Aires |
| chilcas.ar | Laura Yahdjian | Universidad de Buenos Aires |
| cowi.ca | Andrew MacDougall | University of Guelph |
| elliot.us | Elsa Cleland | University of California, San Diego |
| frue.ch | Sabine GÃ¼sewell | ETH Zurich |
| frue.ch | Andy Hector | University of Zurich |
| hall.us | Rebecca McCulley | University of Kentucky |
| hall.us | Jim Nelson | University of Kentucky |
| hart.us | Nicole DeCrappeo | USGS |
| hart.us | David Pyke | USGS |
| hero.uk | Mick Crawley | Imperial College at Silwood Park |
| kbs.us | Lars Brudvig | Michigan State University |
| koffler.ca | Marc Cadotte | University of Toronto Scarborough |
| koffler.ca | Arthur Weiss | University of Toronto Scarborough |
| konz.us | Kimberly Komatsu | University of California, Berkeley |
| konz.us | Melinda Smith | Colorado State University |
| msla.us | Kelly Laflamme | MPG Ranch |
| mtca.au | Suzanne Prober | CSIRO |
| ping.au | Jodi Price | The University of Western Australia |
| ping.au | Rachel Standish | The University of Western Australia |
| potrok.ar | Hector Bahamonde | UNPA - CONICET |
| potrok.ar | Pablo Peri | UNPA - CONICET |
| rook.uk | Mick Crawley | Imperial College at Silwood Park |
| sage.us | Daniel Gruner | University of Maryland |
| sage.us | Louie Yang | University of California, Davis |
| saline.us | Kimberly Komatsu | University of California, Berkeley |
| saline.us | Melinda Smith | Colorado State University |
| sedg.us | Carla D'Antonio | University of California, Santa Barbara |
| sevi.us | Scott Collins | University of New Mexico |
| sevi.us | Laura Ladwig | University of New Mexico |
| sgs.us | Dana Blumenthal | USDA-ARS |
| sgs.us | Cynthia Brown | Colorado State University |
| sgs.us | Julia Klein | Colorado State University |
| sgs.us | Alan Knapp | Colorado State University |
| shps.us | Peter Adler | Utah State University |
| smith.us | Janneke Hille Ris Lambers | University of Washington |
| spin.us | Rebecca McCulley | University of Kentucky |
| spin.us | Jim Nelson | University of Kentucky |
| temple.us | Philip Fay | USDA-ARS |
| trel.us | Andrew Leakey | University of Illinois at Urbana-Champaign |
| ukul.za | Kevin Kirkman | University of KwaZulu-Natal |
| ukul.za | Michelle Tedder | University of KwaZulu-Natal |
| unc.us | Charles Mitchell | University of North Carolina |
| unc.us | Justin Wright | Duke University |
| valm.ch | Martin Schuetz | Swiss Federal Institute for Forest, Snow and Landscape Research |
| yarra.au | Raul Ochoa Hueso | University of Western Sydney |
